# Structure-Based Generation of 3D Small-Molecule Drugs: Are We There Yet?

**DOI:** 10.1101/2024.12.27.629537

**Authors:** Bo Yang, Chijian Xiang, Tongtong Li, Yunong Xu, Jianing Li

## Abstract

Structure-based drug design (SBDD) plays a crucial role in preclinical discovery. Recently, structure-based generative algorithms have been developed to streamline the SBDD process, by generating novel, drug-like molecule designs based on the binding pocket structure of target protein. However, there is no effective metric to evaluate the chemical plausibility of molecules designed by these algorithms, which can limit further applications. In this study, we introduce two new metrics for assessing the chemical plausibility of generated molecules. Along with additional analyses, we demonstrate that these algorithms can generate chemically implausible structures with certain property distributions that differ from those of known drug-like molecules. We also compare results with high-throughput virtual screening hits for three protein targets (the c-SRC kinase, the Smoothened receptor, and the Dopamine D1 receptor). These metrics and analysis methods described here offer valuable tools for model developers and users to assess the chemical plausibility and drug-likeness of generated molecules, ultimately enhancing the use of structure-based generation in drug discovery.

**TOC:** 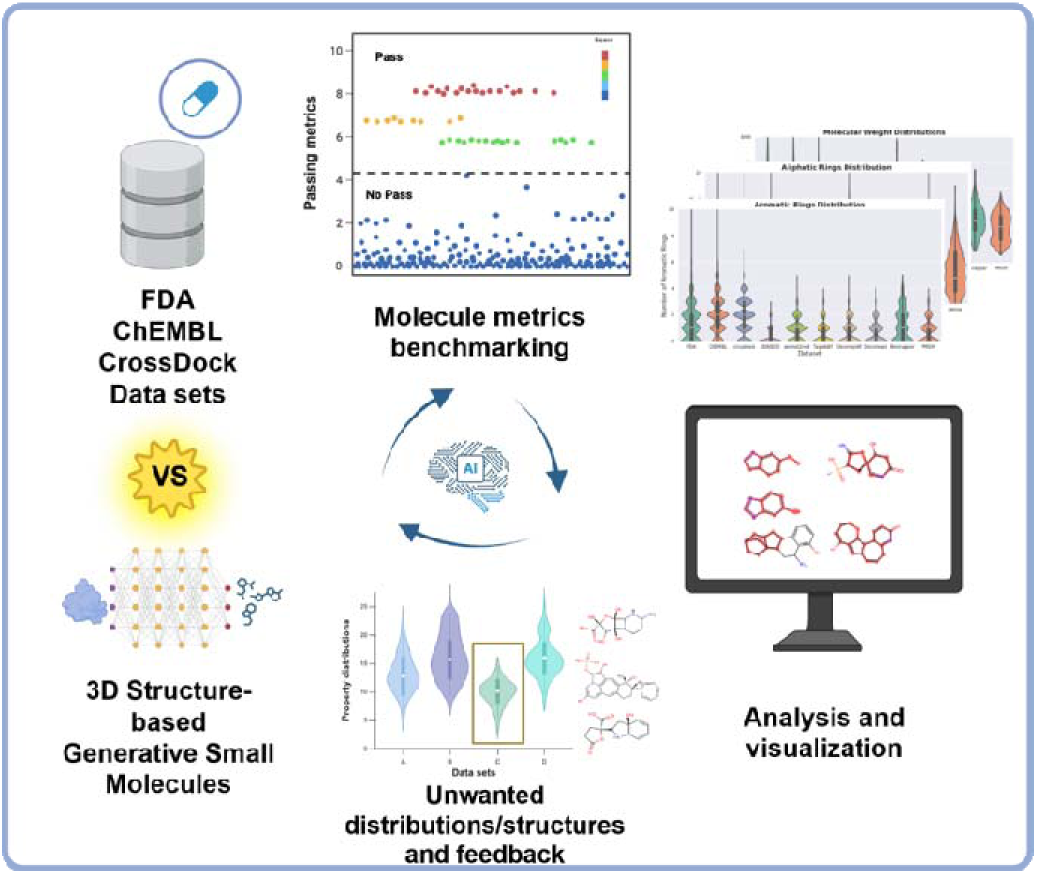

## Introduction

As drug discovery is often costly and lengthy,^1, 2^ numerous efforts have been made to accelerate it, reduce associated costs, and improve the success rate.^3, 4^ Recent advances in artificial intelligence and machine learning (AI/ML) have led to many new applications to enhance drug discovery efforts.^5–7^ Among these, generative AI stands out as one of the most promising. It would be intriguing if generative AI could directly design small-molecule candidates based on information about the therapeutic target, such as the sequence, structure, mechanism, and so forth. A lingering question is: Are we there yet? In other words, can we readily take the AI-designed molecules to any stage of preclinical or even clinical studies? To that end, we developed two novel metrics in this work for evaluating the chemical plausibility of molecules generated based on a three-dimensional (3D) protein target structure. Generally, a structure-based generative model learns the features of protein-ligand interactions during training and generates multiple 3D small molecules expected to bind to a predefined 3D binding pocket (Figure 1).

**Figure 1.**
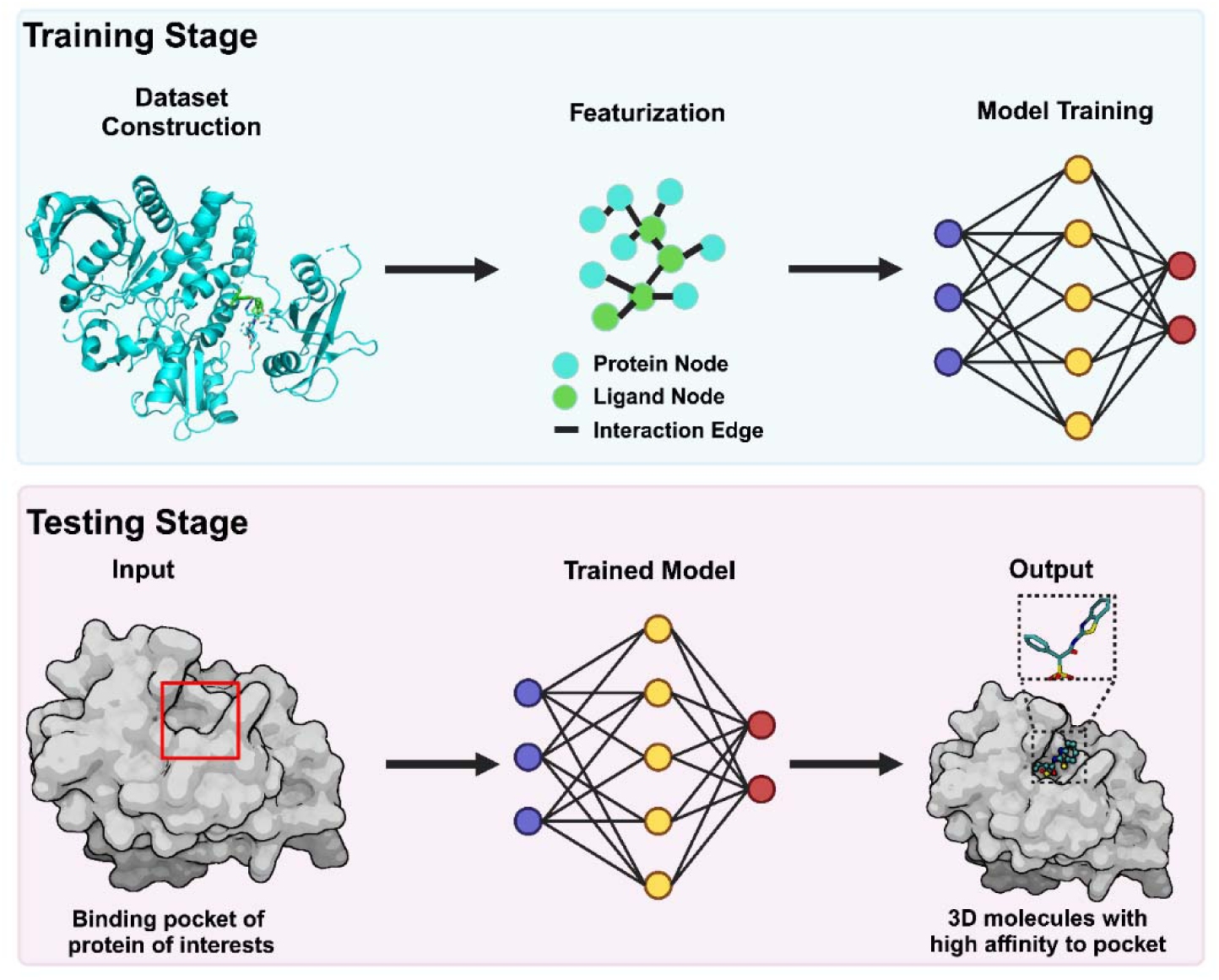
General workflow of structure-based generative algorithms. Protein-ligand complexes are converted into machine-readable features and used as input to train the generative model. Once trained, the model takes the binding pocket of interest as input and generates 3D ligand structures with high affinity for the given binding pocket.

A fundamental idea in drug design is to efficiently search for drug-like candidates from the vast chemical space, estimated to contain over 10^60^ possible compounds.^8^ Medicinal chemists hope to identify compounds that exhibit high potency and specificity for the therapeutic target, as well as good drug-likeness, both in vitro and in vivo, ultimately leading to successful clinical studies. Since the 1990s, the development of structure-based drug design^9^ (SBDD) in tandem with the rapid growth of the Protein Data Bank (PDB) has led to numerous successful discoveries.^10–13^ Currently, a popular SBDD approach is molecular docking to screen large compound libraries, such as the Enamine REAL database^14^ and the ZINC database.^15^ However, most compounds in these libraries are likely to be rejected during docking, primarily due to steric clashes or inadequate interactions with the binding pocket. A more efficient SBDD approach would be to directly “grow” drug-like molecules based on the binding pocket structure (e.g., size, shape, amino acid arrangement, surface interaction, etc.). Built on this idea and powered by the deep learning technique,^16, 17^ structure-based generative algorithms have been developed to learn the features of ligands and interaction patterns in known protein-ligand complex structures and then generate potentially new compounds that mimic these features or patterns.^18–20^ As of spring 2025, there are at least 27k protein-ligand complex structures in the PBD-bind database,^21^ 689k modeled protein-ligand binding complexes in the BindingNet v2 dataset,^22^ and 22.6 million docking protein-ligand complexes in the CrossDocked2020 (hereafter referred to as crossdock) dataset,^23^ which provide fast-growing datasets for training structure-based generative models.

Current structure-based generative models primarily differ in their methods for representing 3D features, incorporating domain knowledge, and generating compound designs (Figure 2). While it was a long-standing challenge to represent the 3D features of protein-ligand complexes, recent breakthroughs include the use of discrete voxelized grids,^24^ continuous methods like 3D molecular graphs and point clouds,^18, 19^ and Euclidean distance metrics.^25^ 3D molecular graphs and point clouds have emerged as the state-of-the-art representations in structure-based generative models because they capture both atomic properties and bond connectivity. By directly encoding precise 3D atomic coordinates, they also incorporate rich geometric information.^18, 26^ These combined features make them more expressive than voxelized grids or distance-based representations for modeling 3D molecules. Furthermore, some generative models incorporate domain knowledge, such as pharmacophores^27^ or explicit protein-ligand interactions,^28^ to guide the generation process. Incorporating domain knowledge may enhance the generation of molecules with better affinity or other properties. However, the restricted generation may also reduce the diversity of generated structures by adding more conditions on the generative process. Finally, two generative strategies have been pursued that can generate molecules auto-regressively (atom-by-atom or fragment-by-fragment) or in one shot.^26, 29^ Overall, almost all the current models were designed to generate molecules conditioned on binding pocket structures with high docking scores. Besides docking scores, commonly used metrics to assess the quality of generated molecules include quantitative estimates of drug-likeness (QED)^30^ and synthetic accessibility (SA) scores,^31^ which are known to have their own limitations.^32, 33^ Recently, Buttenschoen *et al.* introduced *PoseBusters*, a tool designed to evaluate the physical validity of ligand poses generated by deep learning models.^34^ As a rigorous physical tool, *PoseBusters* has driven the development of generative models that produce more physically plausible conformations.^35^ However, while other generative models have been shown to generate chemically implausible structures (i.e., structures that are not chemically stable or synthesizable),^36^ the output of structure-based generative algorithms has yet to be examined rigorously from a medicinal chemistry perspective. Most studies have relied solely on the validity of SMILES strings or 3D structures as criteria for chemical plausibility, supplemented by distribution analyses of 2D structural properties. Nevertheless, these approaches are insufficient for comprehensively evaluating the chemical plausibility.^37–39^ The absence of effective metrics impedes the ongoing development and enhancement of models that generate chemically plausible, drug-like molecules, which we consider essential for practical applications and for advancing drug discovery.

**Figure 2.**
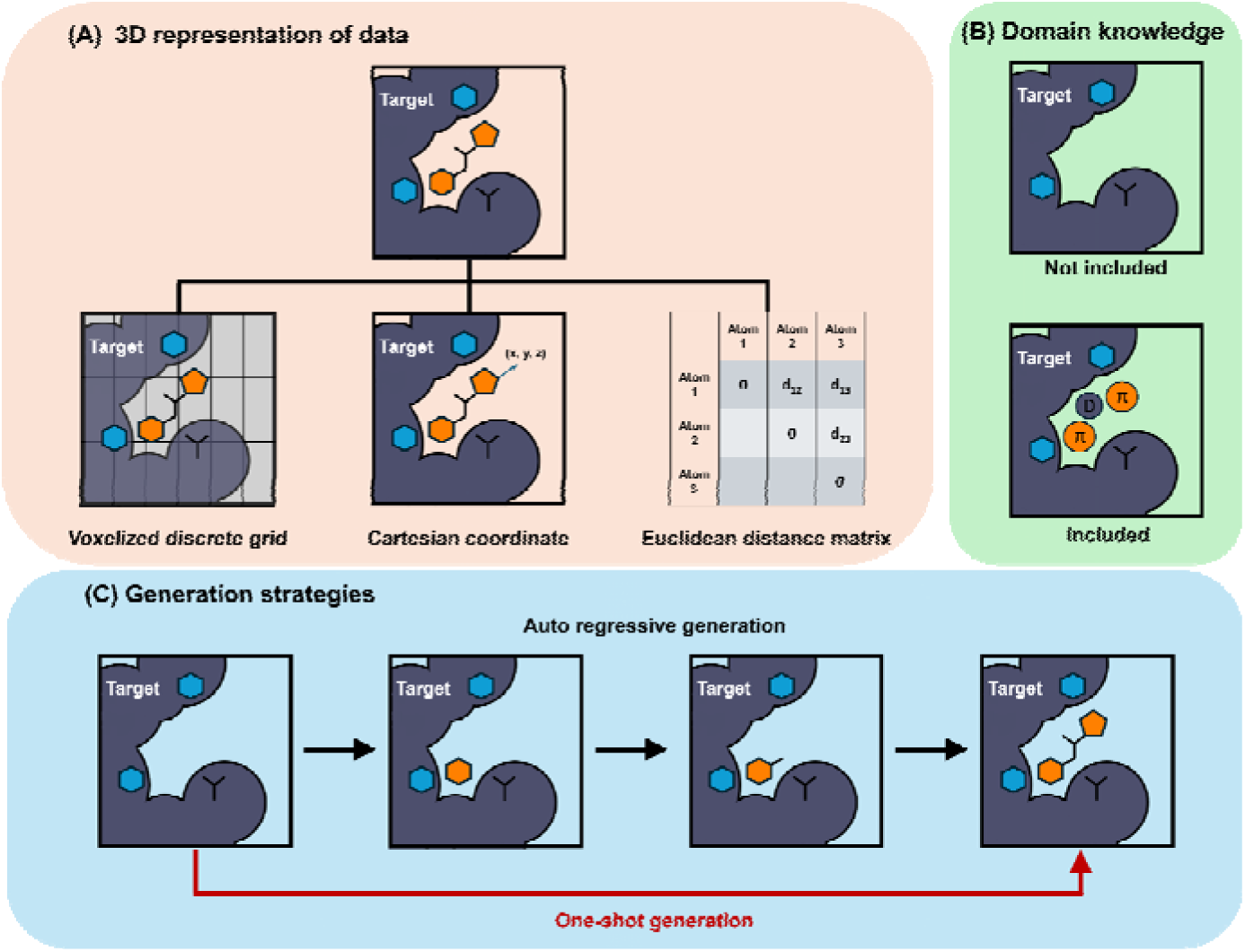
Summary of current 3D structure-based generative algorithms. **(A)** The 3D structures of ligands and proteins can be represented using various formats, including voxelized grids, Cartesian coordinates, and Euclidean distance matrices. More recently, some models represent 3D data as tokens to enable the use of language models for 3D structure-based generation tasks. **(B)** In addition to using protein structures as conditional inputs for molecular generation, domain-specific knowledge such as pharmacophores and protein–ligand interactions can also be incorporated to guide the generative process. **(C)** These models can further be categorized by their generation strategies. Some adopt an auto-regressive approach, generating ligands atom-by-atom or fragment-by-fragment, while others follow a one-shot strategy, such as diffusion models, to generate the full molecule in one step.

To evaluate the ability of generative algorithms to design chemically plausible and drug-like molecules, we introduced two novel metrics developed initially from a survey of drug-like molecules from the ZINC20 and ZINC22 databases.^15, 40^ We applied these, along with an additional metric utilizing ring system occurrence from the ChEMBL database,^36, 41^ to provide a comprehensive assessment of the chemical plausibility of molecules generated by seven structure-based generative algorithms (3D-Generative-SBDD or known as 3DSBDD,^42^ pocket2mol,^19^ TargetDiff,^18^ DecompDiff,^43^ PMDM,^39^ Decompopt,^38^ and MolSnapper^35^), in comparison with three control groups (FDA-approved small molecule drugs, ChEMBL clinical small molecules, as well as the crossdock dataset^23^) that are believed to contain drug-like molecules. Overall, we found that molecules produced by the generative models are less drug-like compared to those from the control groups, largely due to insufficient chemical plausibility and drug-likeness. Further property-distribution analysis and visualizations identify contributory factors—possibly stemming from the training datasets and model architectures—that likely underlie this performance gap. To reduce potential bias from the target distribution of the crossdock test set, we carefully chose three representative protein targets—the c-Src kinase (PDB ID: 7WF5),^44^ the Smoothened receptor (PDB ID: 5L7I),^45^ and the Dopamine D1 receptor (PDB ID: 7CKY)^46^—and compared the generated molecules with high-throughput virtual screening (HTVS) hits. The generative algorithms could design molecules with high predicted binding affinity, but they consistently faced challenges in generating chemically plausible and drug-like compounds in our studies. Building on these findings, our metrics and analyses offer practical guidance for refining structure-based generative algorithms to yield more chemically plausible, drug-like molecule designs.

## Results

### Are all the generated molecules chemically plausible and drug-like?

To ensure a fair comparison of structure-based generative models, all selected generative models were trained and tested on the same subsets of the CrossDock2020 dataset.^23, 42^ To avoid training bias, we utilized pre-trained models to generate 100 molecules for each of the 100 proteins in the test set, aiming for a total of 10,000 molecules for the benchmarking study. However, we found that not all algorithms generated exactly 10,000 molecules (presumably due to their algorithm design, see a summary in Table S2). Some generated molecules were duplicates or invalid, as indicated by a ‘Non’ value returned during the conversion from SMILES or structure data file (.sdf) to a Mol object using the RDKit package.^47^ Table S2 also summarizes the generation efficiency for each algorithm and the device used for generation for reference, which is generally consistent with earlier studies.^48^

With these molecular structures generated by different models, we applied several metrics to assess their chemical plausibility. The first metric involves extracting ring systems from each generated molecule and recording their occurrence in the ChEMBL database.^36, 41^ ChEMBL is a manually curated database of bioactive molecules with drug-like properties. If a ring system is uncommon in the ChEMBL compounds, its chance of being either chemically plausible or drug-like can be low. The second and third metrics evaluate whether the Bemis-Murcko^49^ (BM) scaffolds of the molecules appear among drug-like molecules in the ZINC20 (870 million) and ZINC22 (54.6 billion) databases. If a molecule’s BM scaffold isn’t present in these databases, it might suggest instability or a lack of drug-like properties. Additionally, it is worth noting that the “drug-like molecules” in the ZINC databases are predicted based on molecular properties and may not necessarily be truly drug-like, while some ZINC compounds have not been synthesized but are predicted to be synthesizable. Additionally, a BM scaffold can be absent from ZINC while still being chemically plausible, and the same is true for the ring systems in ChEMBL. Our results clearly indicate that the three metrics used in this work show high agreement between the three control groups (Figure 3), while inconsistency is observed among the generated compounds.

**Figure 3.**
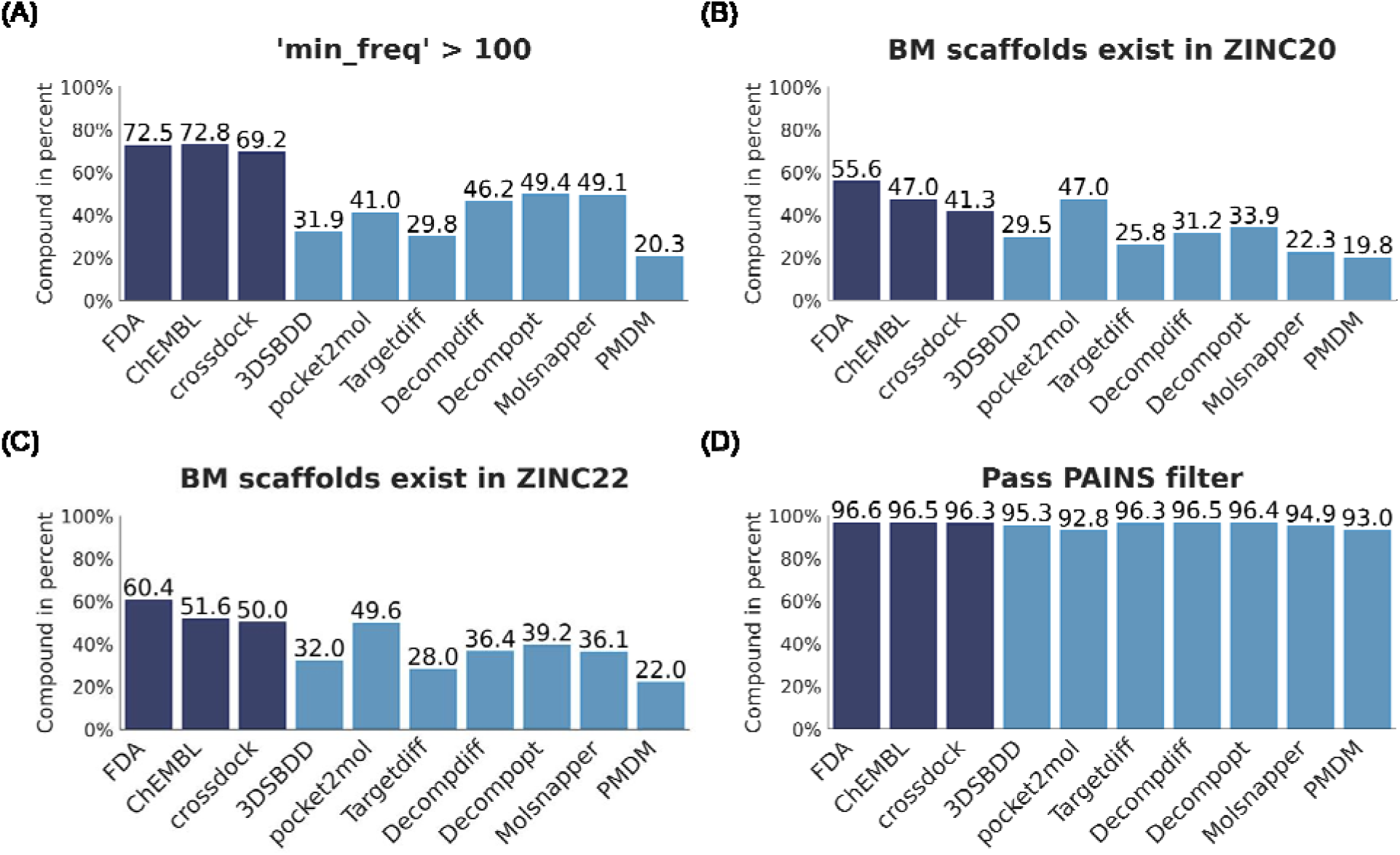
Percentage of compounds that are evaluated with three metrics and the PAINS filter. All values are normalized. **(A)** The “min_freq > 100” metric is calculated by extracting the ring systems in each molecule and documenting the frequency of each ring system’s occurrence in the ChEMBL database. If the least frequent ring system (some molecules may contain multiple ring systems) appears fewer than 100 times in ChEMBL, the molecule is considered potentially unstable or non-drug-like. The ratio is determined by dividing the number of molecules that meet this criterion by the number of unique molecules containing ring systems. **(B-C)** “BM scaffolds” are calculated by extracting the scaffolds of drug-like molecules from ZINC20 and ZINC22 databases. If the BM scaffolds of generated molecules are not found in these databases, it may suggest that the generated structures are either complex to synthesize or chemically implausible. The ratio is calculated by dividing the number of BM scaffolds present in ZINC by the total number of BM scaffolds from the generated molecules. **(D)** Percentages of molecules that pass the PAINS filter.

The three control groups (labeled as FDA, ChEMBL, and crossdock) consistently perform well on all metrics, which suggests that compounds in these groups are likely to share similar properties or drug-likeness. Over 69% of control-group molecules pass the ChEMBL ring-system metric, compared to fewer than 50% of algorithm-generated molecules (Figure 3A). A consistent difference was also found between our control molecules and the generated molecules using the two ZINC-based metrics, which measure the percentage of BM scaffolds from ZINC20 and ZINC22 compounds (Figures 3B and 3C, respectively). The three control groups achieve about 10% higher results in both metrics than the generated groups, except for pocket2mol. Moreover, the highest percentages were often observed with the FDA-approved small molecules, followed by the ChEMBL clinical compound set and the crossdock set, indicating these metrics can identify the most druglike data set. This distinction between FDA-approved drugs and the other two control sets suggests that the presence of BM scaffolds may serve as a strict indicator of drug-likeness. Compared to molecules from the control groups, those generated by algorithms may be less chemically plausible or less drug-like in general, as revealed by the metrics applied here. In the later sections of this work, we analyzed the chemically implausible structures in the generative design and provided examples and additional analysis to support our findings.

Notably, the filter for pan-assay interference compounds, PAINS,^50^ did not effectively exclude chemically implausible molecules, as shown by over 92% of molecules passing the filter across all datasets (Figures 3D and S4). The PAINS filter, along with other commonly used filters in our tests (Figure S4), was designed to identify chemical compounds that contain the substructures often linked to false-positive results in high-throughput screening experiments. These filters, which rely on substructure libraries, cannot verify the chemical stability or viability of newly generated molecules. It is possible that some of the generative models already account for PAINS. Overall, the PAINS filter and other filters may have subtle effects in removing chemically implausible molecules during generative design.

### What can cause chemical implausibility and low drug-likeness in generative design?

It is impractical to visually inspect our generated molecules one by one. To better understand why structure-based generative models design implausible compounds, we first plotted the distributions of several structural properties and descriptors and compared them with control datasets. Overall, the control groups exhibited similar distributions, indicated by the radar charts (Figure 4). In contrast, the distributions of molecules generated by algorithms differ substantially from those of the control groups, particularly with respect to chiral atom count, aliphatic ring count, and aromatic ring count.(Figure 4). Detailed distributions, shown as violin plots, also highlight the agreement within the control group and the discrepancies between the controls and the generated molecules (Figure S2). To better understand these discrepancies, we examined some of the implausible designs and identified possible causes for them from a medicinal chemistry perspective.

**Figure 4.**
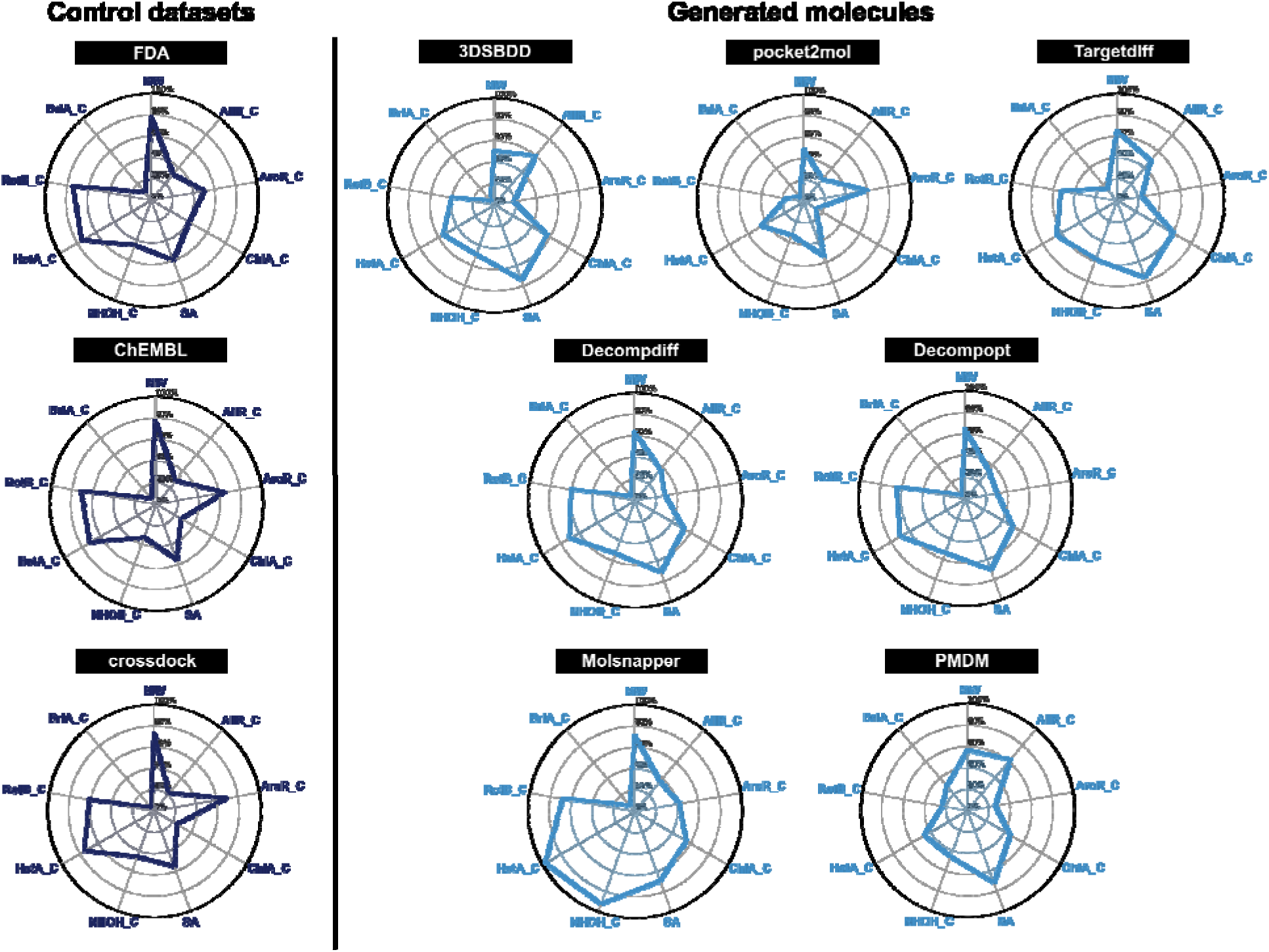
Radar charts to compare chemical property distributions in our datasets. The parameters we examined are defined as follows. *MW:* molecular weight; *AliR_C:* aliphatic ring count; *AroR_C:* aromatic ring count; *ChiA_C:* chiral atom count; *SA:* synthesis accessibility; *NHOH_C:* NH/OH group count; *HetA_C:* heteroatom count; *RotB_C:* rotatable bond count; *BriA_C:* bridgehead atom count. Percentages were calculated using the following values as denominators, with the average value serving as the numerator: MW=500, AliR_C=4, AroR_C=3, ChiA_C=6, SA=6, NHOH_C=6, HetA_C=10, RotB_C=8, and BriA_C=2.

First, chemically unstable structures or substructures in the designs present a common issue, even in low molecular-weight designs. For example, 3DSBDD and pocket2mol generated relatively smaller molecules than the others. The average molecular weight of molecules from these methods is under 255 Da, compared to over 280 Da for molecules from other algorithms tested in this work (Figure 5A). While these designs may have some advantages regarding drug-likeness parameters or metrics (such as synthesizability), the very restricted number of atoms and feasible bonds likely requires more sophisticated designs. Nevertheless, we identified several unreasonable ones in the generated datasets, such as a sp-hybridized carbon atom in a five-member ring (Figure 5B). Chemically unstable structures can be found in every generated dataset, suggesting that some fundamental chemical rules may still be missing in current algorithms or models.

**Figure 5.**
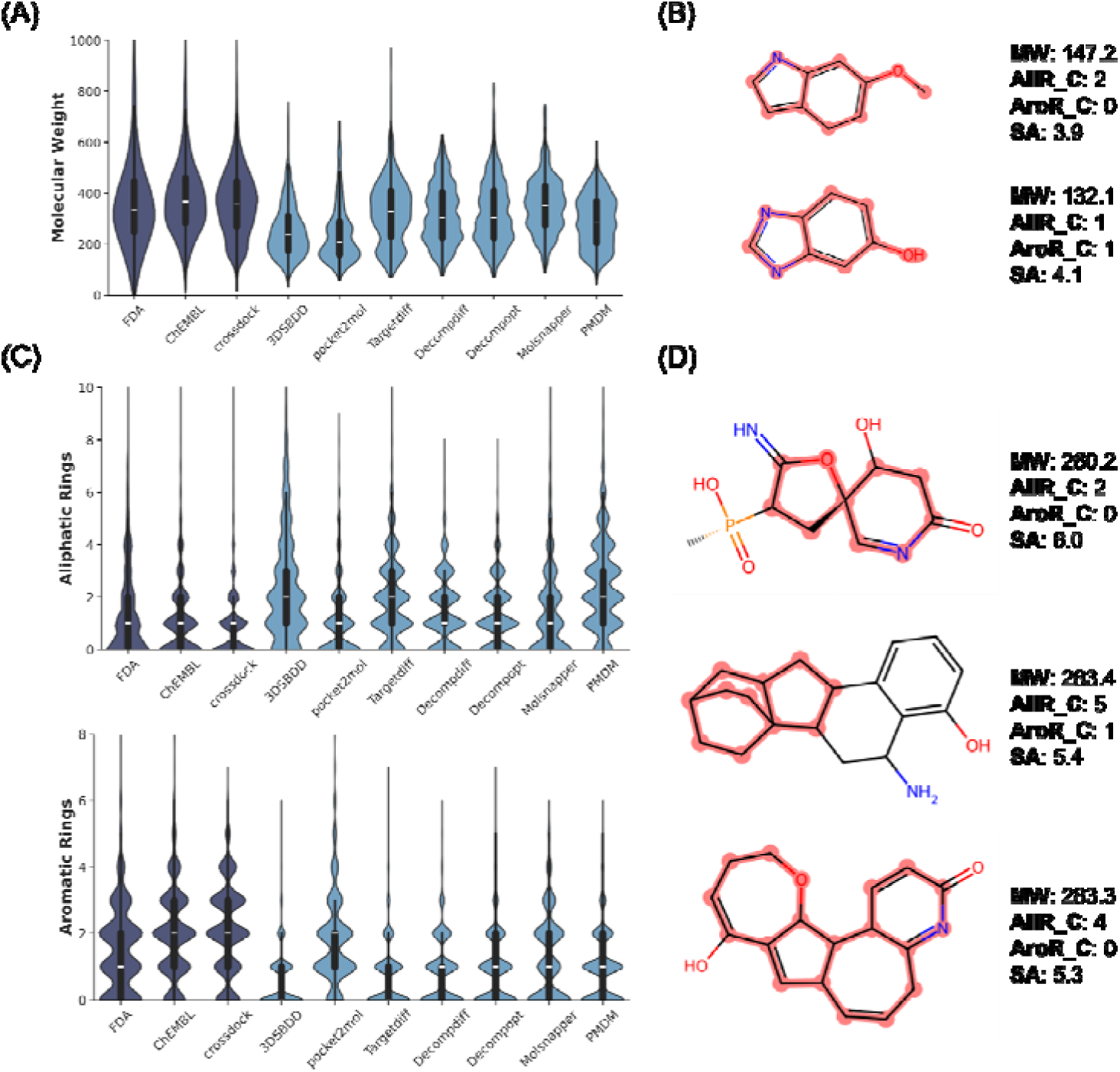
Chemical property distributions show differences between the generated molecules and control datasets. Corresponding chemically implausible structures are selected from molecules that didn’t pass ChEMBL ring metrics, and substructures associated with the distributions are highlighted in red by RDKit. **(A)** The molecular weight distribution across datasets. **(B)** Examples of molecules with low molecular weight, generated by 3DSBDD or pocket2mol. **(C)** Number of aliphatic rings and aromatic rings distributions across datasets. **(D)** Example of chemically implausible molecules with multiple aliphatic rings and one or no aromatic rings.

Second, the generated molecules typically contain too few aromatic rings but too many aliphatic ones. The average number of aromatic rings per generated molecule for each algorithm ranged from 0.5 to 1.75, with most algorithms generating fewer than 1 aromatic ring per molecule. In contrast, the control groups typically averaged more than 1.5 aromatic rings per molecule, with ChEMBL and crossdock molecules having values close to 2 (Figures 5C and S6). Further, aside from pocket2mol, the other algorithms produced more aliphatic rings, with average counts per molecule ranging from 1.35 to 2.5. This is generally higher than the control groups, which have fewer than 1.3 aliphatic rings per molecule on average (Figure 5C). There can be a bias toward aliphatic ring systems over aromatic ones in these generative algorithms. Several examples are shown in Figure 5D to illustrate this bias in actual designs. Such bias can allow the designed molecules to adopt non-flat scaffolds and maximize the contacts at the binding pocket, but it may also lead to challenging synthesis or chemically implausible structures.

Third, the generated molecules typically have more chiral centers. On average, molecules generated by each algorithm contain between 2.7 and 3.6 chiral atoms per molecule, with most algorithms producing more than 3.0. In comparison, crossdock and ChEMBL molecules average around 1.6 chiral centers, while FDA molecules average approximately 2.5. Current algorithms tend to generate more structurally complex molecules, which may be harder to synthesize (Figures 6A and S6). Indeed, we observed higher synthetic accessibility scores (predicted to be more challenging to synthesize^51^) for the generated molecules, with an average SA score of over 4.1 (Figure 6B). Some typical designs with multiple chiral centers in complex ring systems are shown in Figure 6C. However, the three control groups show lower SA scores: 3.6 for FDA-approved small-molecule drugs, 3.3 for ChEMBL clinical small molecules, and 3.3 for ligands in the crossdock dataset. Moreover, we observed that the distribution of molecules produced by the algorithms differs significantly from that of their training set, crossdock (Figure 4). This suggests that additional effort is required to learn the chemical prior information encoded in the training molecules.

**Figure 6.**
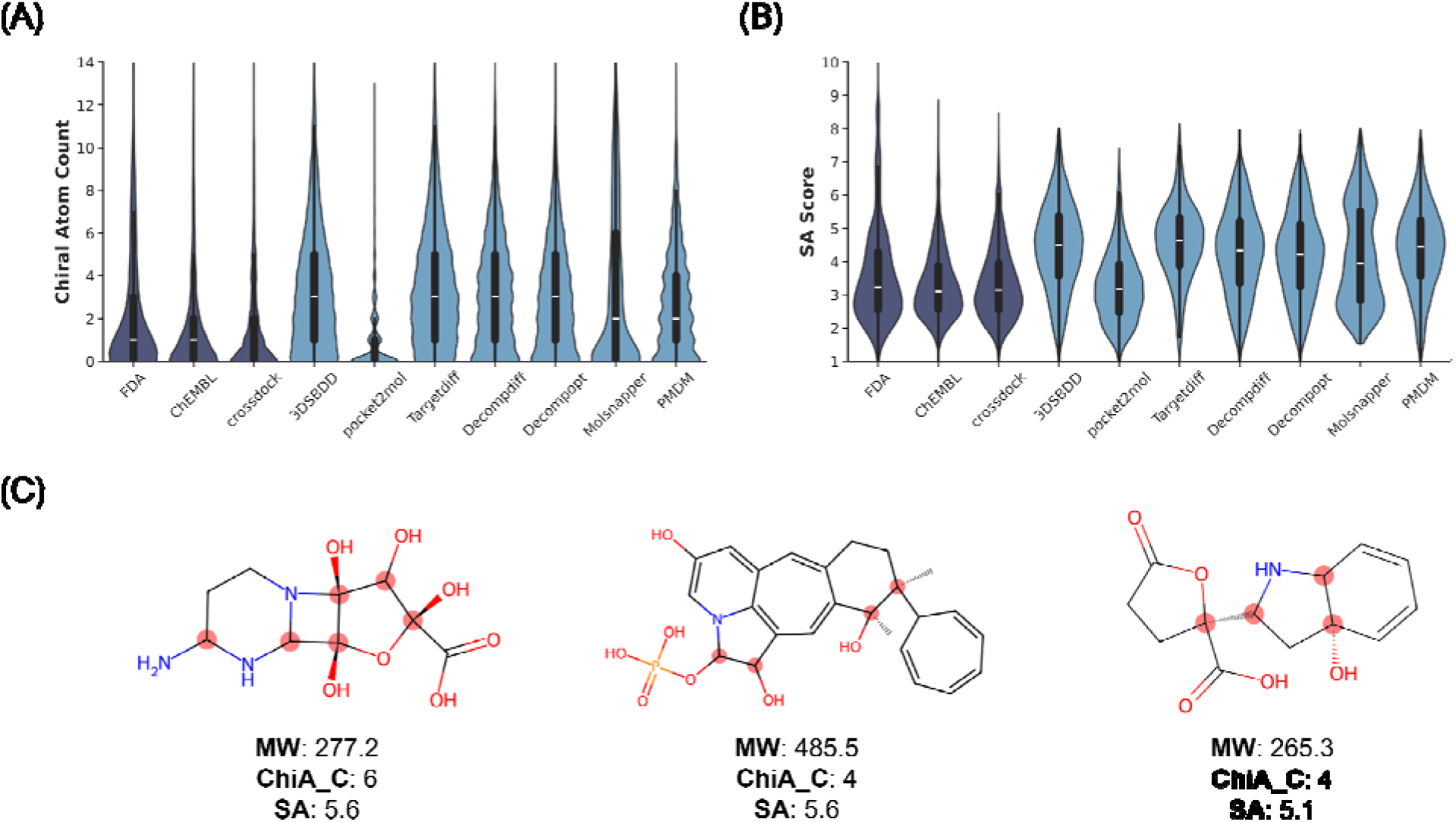
Distributions associated with structural complexity show differences between generated molecules and control datasets. Corresponding chemically implausible structures are selected from molecules that didn’t pass ChEMBL ring metrics, and substructures associated with the distributions are highlighted in red by RDKit. **(A)** Chiral atom count and synthetic accessibility (SA) score distributions across datasets. **(B)** Example of molecules with multiple chiral atoms.

Finally, generated molecules have more NH and OH groups compared to the control groups (Figures 7A and S6). On average, algorithm-generated molecules contain 2.8 to 5.5 NH/OH groups per molecule, whereas the control groups average fewer than 2.8. We highlight several example molecules (Figure 7B) generated by the algorithms that contain multiple NH and OH groups. Although NH/OH groups are common in drug-like molecules, an excess of these groups may negatively impact permeability and other pharmacokinetic (PK) properties. The Lipinski’s Rule of Five suggests that a molecule with more than 5 hydrogen bond donors (OH and NH groups) may have poor oral bioavailability, as they can increase molecular polarity and reduce membrane permeability. Excessive NH and OH groups may hinder the drug’s ability to cross cell membranes and reach its target and introduce additional metabolically labile sites on the molecule. The abundance of OH/NH groups may suggest that these algorithms have learned favorable interactions between the molecule and protein, as OH/NH groups significantly contribute to binding through hydrogen bonding and electrostatic interactions. However, drug-likeness and chemical insights are not well captured due to the structures they produced.

**Figure 7.**
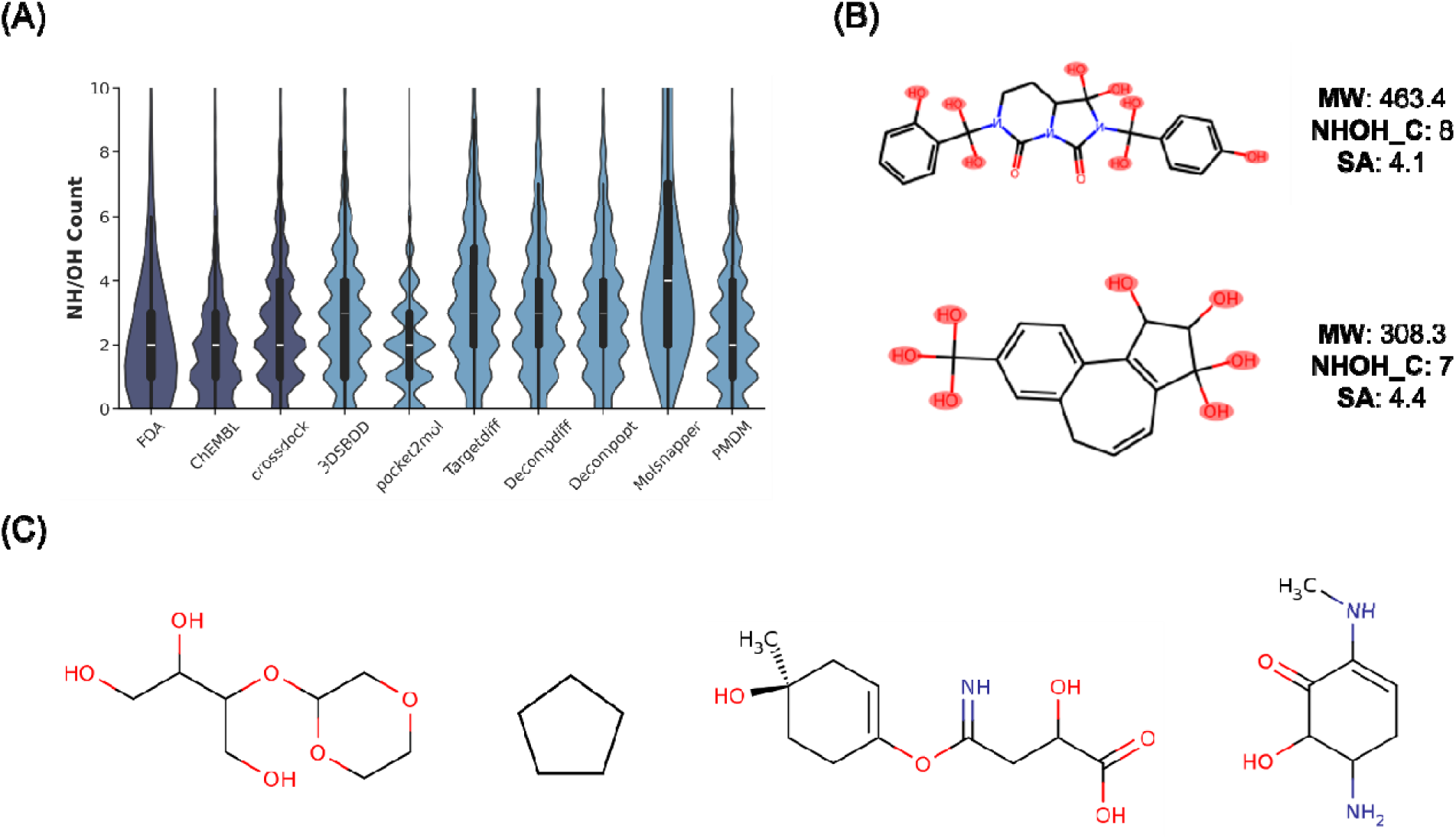
Specific functional groups distribution and example molecules. **(A)** The distribution of NH and OH group count across datasets reveals that the algorithm prefers generating NH and OH groups; substructures associated with the distributions are highlighted in red by RDKit. **(B)** Example of molecules with multiple OH/NH groups. **(C)** Four molecules that could meet all three metrics we applied, but are still not drug-like or difficult to synthesize.

### How does generative design compare with high-throughput virtual screening (HTVS)?

As HTVS and the underlying technology, molecular docking, have been widely used, we assessed the HTVS and generative design approaches using the same protein targets. Notably, most of our tested algorithms were trained on the crossdock dataset; however, the protein targets in that dataset do not adequately represent the distribution of therapeutic targets of FDA-approved small molecules or ChEMBL clinical compounds. For instance, no G protein–coupled receptors (GPCRs) are included in the crossdock training and test set (Table S3). The variation in protein targets included in the training set may raise questions about the fundamental differences between the generated molecules and actual drugs or drug-like molecules, as well as how accurate these algorithms are for new targets, especially compared to HTVS or docking. While a thorough comparison might require experimental validation, in this work, we compared the generated molecules with the top-scoring molecules from HTVS, mainly focusing on the docking scores and three metrics. To ensure a fair comparison, we selected three drug targets: the C-Src kinase (PDB ID: 7WF5),^44^ the Smoothened receptor (PDB ID: 5L7I),^45^ and the Dopamine D1 receptor (PDB ID: 7CKY).^46^ They were chosen because they were not in the training or test datasets used by these generative algorithms, their ligand-binding sites are deep and well-defined, and no conserved water molecules are present in the binding sites.

We only tested 3DSBDD, pocket2mol, and MolSnapper to generate 500 potential ligands for each of the three protein targets because DecompDiff and DecompOpt currently do not support user-defined proteins, and PMDM and TargetDiff failed to generate enough designs. Particularly, MolSnapper was evaluated under two conditions—generation with and without pharmacophore information derived from crystal ligands—to enable a more thorough comparison. In parallel, we carried out HTVS with one-tenth of the Enamine HTS collection (∼174,000 molecules). The average docking scores for all ligands from Enamine HTS collection are -8.3, -8.9, and - 8.0 for docking to the PDB structures 7WF5, 5L7I, and 7CKY, respectively. The top 500 molecules with the lowest Autodock Vina docking scores from HTVS were identified as HTVS hits. The average docking scores of HTVS hits are -12.2, -12.4, and -10.9 for 7WF5, 5L7I, and 7CKY, respectively, which are better compared to those of generated molecules (Figure 8A). Overall, the generated designs from each algorithm show more favorable docking scores (more negative) than the average of all ligands but not the top 500 HTVS hits (Figure 8A). While over 75% of HTVS hits and under 65% of generated designs passed the ‘min_freq > 100’ measure, fewer than 55% of these molecules from either approach have passed the BM scaffolds metrics (Figures 8B-D). Since the BM-scaffold-based metrics are likely more stringent than the ‘min_freq > 100’ one, our results suggest that the molecules generated for these three targets are less drug-like than the training dataset (crossdock). Their chemical plausibility and drug-likeness in general were lower than HTVS hits from the Enamine HTS collection. These results highlight the importance of including chemical plausibility assessments in evaluating generative models to ensure high docking scores correspond to practically chemically plausible, drug-like molecules.

**Figure 8.**
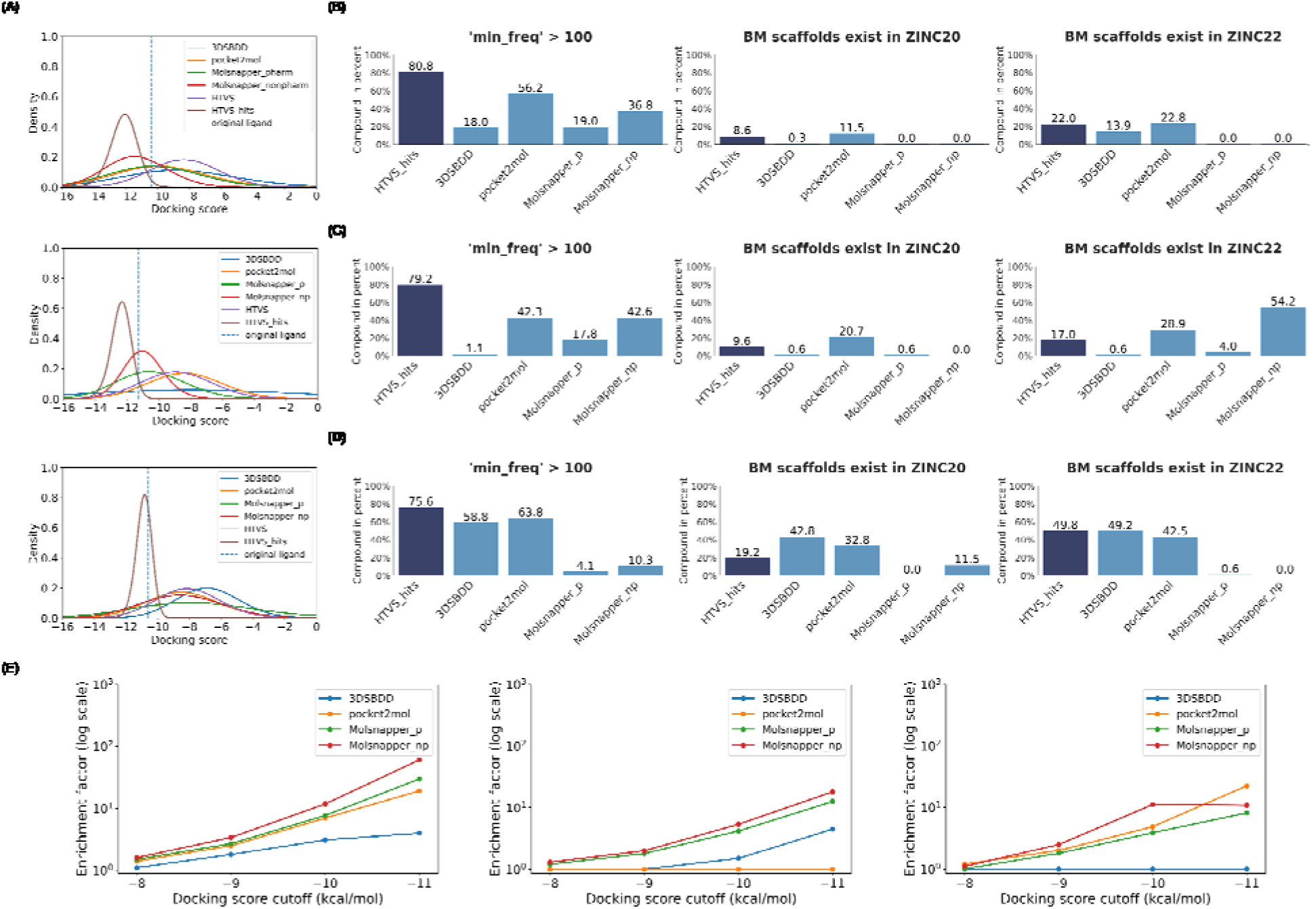
Comparison of molecules generated by different algorithms with high-throughput virtual screening (HTVS) hits against C-Src kinase (PDB ID: 7WF5), the Smoothened receptor (PDB ID: 5L7I), and the dopamine D1 receptor (PDB ID: 7CKY). For MolSnapper, “p” indicates inclusion of pharmacophore information from the crystal ligand during generation, whereas “np” indicates that pharmacophore information was not used. Rows correspond to protein targets: first row, 7WF5; second row, 5L7I; third row, 7CKY. (A) Vina score distributions for generated molecules, one-tenth of the Enamine HTS collection, and HTVS hits. (B-D) Percentages of compounds satisfying three quality-control metrics: ring_freq > 100; BM scaffolds exist in ZINC20; and BM scaffolds exist in ZINC22. For this analysis, the top 500 HTS molecules ranked by Vina score were used to calculate pass rates. (E) Enrichment factors for Vina scores of generated molecules relative to one-tenth of the Enamine HTS collection (left to right: 7WF5, 5L7I, 7CKY).

We also compared HTVS and generative design in terms of the balance between accuracy and efficiency, using the enrichment factor (EF)^48^ and the time to generate a compound. The EF was defined as the ratio of the generated hit rate to the HTVS hit rate, with a given docking score threshold to determine a hit. We calculated the EF with a threshold varying from -8 kcal/mol to -12 kcal/mol, and a value of EF greater than 1 may indicate the effectiveness of generative design (Figure 8E, Table S4). We observed a general trend of EF increasing as the threshold becomes lower or more negative. MolSnapper consistently showed EF above 1, with or without pharmacophore information, but 3DSBDD and pocket2mol failed to generate molecules with strong docking scores for at least one target, indicated by EF close to 0 for at least one target (Figure 8E, Table S4). Given that the average docking time per molecule is 3.37 seconds for Vina-GPU 2.1,^52^ and that all algorithms generated one molecule in under 30 seconds on average (Table S2). We suggest that with EF greater than 10, generative design with a robust algorithm like MolSnapper may provide a time-efficient approach for identifying hit molecules. However, in the three protein targets we tested, high EF was only achieved with a relatively high threshold (below -10 kcal/mol) and was highly dependent on the target.

## Discussion

In summary, we have identified chemical implausibility in AI-designed molecules and several shortcomings of current generative algorithms for future enhancement. Two effective metrics based on the occurrence of BM scaffolds in ZINC20 and ZINC22 were developed to evaluate the generated designs. Along with the ring occurrence metric in ChEMBL, these metrics show that some generated molecules are neither chemically plausible nor drug-like when compared to the three control groups. Further structural and property analyses, supported by visual inspection, highlight potential weaknesses in the generated molecules that may contribute to their chemical implausibility and lower drug-likeness compared to the controls. Consistent with these findings, case studies on the c-Src kinase, the Smoothened receptor, and the Dopamine D1 receptor showed that, although generative algorithms were able to produce molecules with favorable docking scores, their chemical plausibility and drug-likeness are generally lower than HTVS hits from a chemical library like the Enamine HTS collection. These results underscore the necessity of incorporating chemical plausibility assessments into generative model evaluation to ensure that high docking scores translate into practically chemical plausible, drug-like molecules.

Beyond chemical plausibility and drug-likeness, which can be assessed from the 2D molecular graph, the quality of 3D binding poses of generated molecules is equally critical. Several studies have reported that current generative models often produce molecules with unreasonable 3D conformations, characterized by high strain energies and spatial clashes.^37, 53^ Using PoseBuster,^34^ PoseCheck^54^ and Torsion library,^55^ we evaluated the 3D binding poses of molecules generated by each algorithm on the crossdock test set and obtained results consistent with these previous studies (Table S5). Compared with ligands from the crossdock test set—whose binding poses were generated by docking—most of the algorithm generated conformations performed significantly worse on these metrics (Table S5). Post-processing strategies such as energy minimization and redocking are known to alleviate some of these issues;^37, 53, 56^ however, even though this was not the primary focus of our study, the quality of 3D conformation generation remains an important challenge for 3D structure–based generative models.

It is evident that current 3D structure–based generative models exhibit suboptimal performance in capturing chemical prior information from the training set, as reflected by the chemically implausible structures generated, the low pass rates of our proposed metrics, and the low retrosynthesis success rates predicted by SynFormer^57^ compared with control groups (Table S6). Commonly used metrics such as QED and SA fail to fully capture this lack of chemical plausibility (Table S6). Several strategies may help address this limitation. A straightforward approach is to apply compound filters,^41, 58^ visual inspection, or carefully designed screening workflows^36, 59^ to remove chemically implausible or non-drug-like molecules. Another approach is to pretrain models on synthesized or realistic molecular datasets to embed stronger chemical knowledge prior to engaging in structure-based generative tasks. Such pretraining is relatively common in chemical language models^60–62^ but remains underexplored for graph-based models. Alternatively, reinforcement learning guided by chemical plausibility metrics—such as those introduced in this study—could steer molecular generation toward more plausible and drug-like structures.^63, 64^ In addition, fragment-based generation approaches^65, 66^ or synthesis-accessibility-aware models^67^ can improve plausibility by leveraging the prior information encoded in fragments, building blocks, or reaction libraries. However, the breadth of chemical space explored is inherently constrained by the predefined nature of these resources. Finally, several recent studies have introduced AI-based post-processing models capable of either repairing problematic structures^68^ or generating structurally similar but valid alternatives^57^, which may provide another effective route for mitigating these challenges.

Another factor that may contribute to the suboptimal performance of those models on both 2D and 3D prospectives is the quality of the crossdock dataset. As its name suggests, crossdock was constructed by docking ligands from known protein-ligand complexes into protein with similar binding pockets. Thus, the protein-ligand complexes used for training and testing may not represent optimal binding pairs. As shown in Figure S8B, the van der Waals (vdw) volumes of ligands in the test set exhibit only a weak correlation with the pocket volume of the proteins, which may indicate the suboptimal interactions due to the crossdocking nature. Meanwhile, the crossdock dataset does not adequately represent the diversity of therapeutic targets. For example, G protein-coupled receptors (GPCRs) are absent from both the training and test set (Table S3). This bias could influence its performance on more comprehensive, real-world drug discovery cases. Given that crossdock is currently the most widely used training set for 3D structure-based generative models, adopting newer and more comprehensive datasets — such as BindingNetV2^22^ and Large Scale Docking (LSD) database^69^— could improve both performance and generalization by providing more robust and realistic protein-ligand complexes.

## Conclusion Remarks

Although challenges remain, we still believe that 3D structure-based generative models hold tremendous potential to accelerate preclinical drug discovery. By leveraging structural information, these models can design diverse classes of pharmaceutical molecules,^70, 71^ explore novel chemical space, generate molecules with a favorable Vina docking score, and provide valuable structural insights for medicinal chemists. Although these algorithms currently generate both promising and misleading molecules, further development is expected to produce higher-quality candidates, ultimately encouraging their routine use in medicinal chemistry. Together, our metrics and the analyses presented here provide valuable insights for model developers to improve these algorithms and enhance their utility in drug discovery. The code and data used in the work can be found at https://github.com/Kartinaa/Benchmarking_gene_model.

## Method

### Data curation

FDA approved small molecule drugs control set was extracted from DrugBank 5.1.12 database. After downloading the full database, a dedicated python script was used to extract the approved small molecule drugs as InChI format and save locally. The ChEMBL clinical small molecules control set was downloaded from the ChEMBL 34 database using website filters to exclude approved small molecules, ensuring no redundancy with the FDA-approved control set. The CrossDocked2020 control dataset was created by converting all 3D ligands from the CrossDocked2020 dataset into SMILES format and removing duplicates. All control datasets are made publicly available at https://github.com/Kartinaa/Benchmarking_gene_model.

Regarding the datasets used in the metrics: ZINC20 drug-like molecules (0.87 billion) were downloaded from the ZINC20 database’s drug-like subset with the “Wait OK” label. ZINC22 drug-like molecules (54.6 billion) were downloaded from the drug-like subsets in 2D trenches from the ZINC22 database. It is important to note that many of the synthesis-on-demand molecules in ZINC have not yet been synthesized. Therefore, their synthesizability is not guaranteed. For reference, the ZINC20 drug-like subset contains 688 FDA-approved small-molecule drugs, 2,621 ChEMBL clinical compounds, and 1,080 molecules from CrossDock2020. In comparison, the ZINC22 drug-like subset contains 810, 1,560, and 1,669 molecules in these control sets, respectively.

### Molecule generation

As outlined in the manuscript, all algorithms compared in this study use the same training and test sets. The goal was to generate 100 molecules for each protein in the test set using pre-trained models, theoretically resulting in a total of 10,000 molecules per algorithm (100 molecules × 100 proteins in the test set). However, most algorithms did not generate exactly 10,000 molecules, and ensuring precisely 100 molecules per test protein was challenging. Since this study focuses on evaluating the chemical plausibility of the generated structures, the reasons for this variability are not discussed further.

All molecule generations were conducted using the default configurations or those recommended by the respective authors. One critical factor in molecular generation is the number of heavy atoms in the generated molecules, as this parameter can substantially influence model performance. The strategies used by each algorithm to determine the number of heavy atoms are summarized below. (1) Test set proteins. 3DSBDD and pocket2mol employed a frontier atom prediction model to decide whether to add additional atoms, thereby predicting the total number of heavy atoms. PMDM, DecompDiff, and DecompOpt used the heavy atom count of the corresponding ligands from the test set proteins. TargetDiff determined the heavy atom count based on a predefined distribution derived from pocket size. MolSnapper used the ground-truth heavy atom number as the mean value to sample from a predefined distribution. (2) User-defined proteins. 3DSBDD and pocket2mol again predicted the heavy atom count for each generated molecule. PMDM used a user-defined heavy atom number. MolSnapper used the user-defined value as the mean for sampling from a predefined distribution. TargetDiff applied the same pocket-size–based predefined distribution as in the test set. DecompDiff and DecompOpt could only operate on the test set and were therefore excluded from user-defined target generation.

To ensure reproducibility, we briefly outline the parameters used for several algorithms: For Molsnapper, the ‘num_pharma_atoms’ flag was set to ‘20’ and ‘clash_rat’ was set to 0.1, as indicated in the GitHub repository. For Decompdiff, the ‘prior_mod’ flag was set to ‘ref_prior’ following configuration file. For Decompopt, the ‘dup_num’ variable in ‘sample_compose.sh’ was set to 0, avoiding re-docking molecules into the binding pocket to improve generation efficiency. While this setting may reduce the performance of generated molecules on docking-based metrics, we expect that it does not significantly impact the chemical plausibility of the generated structures.

### Metrics incorporation and property calculations

Before processing and evaluating the generated molecules, all structures were standardized using MolVS.^72^ Specifically, salts were removed, tautomers were standardized, charges were neutralized, isotopes were standardized, functional groups such as nitro groups were normalized, and the molecules were sanitized. The canonical SMILES of standardized molecules generated by each algorithm are saved for further use. For ring system-based metrics, ring systems for each molecule were extracted using ‘useful_rdkit_utils’ python library, which incorporates the ring system frequency data from the ChEMBL database. For the ZINC20 and ZINC22 drug-like sets, ring system frequencies were calculated using the same Python library and served as baselines for comparison with the ring system frequencies of the generated molecules. For BM scaffold-based metrics, the BM scaffolds were extracted from the ZINC20 and ZINC22 drug-like sets using the RDKit Python library. After removing redundancies, the BM scaffolds from each dataset were saved locally and used as baselines to evaluate the occurrences of BM scaffolds in the generated molecules. All chemical properties were calculated using the RDKit Python library, and visualizations were created using Matplotlib, Seaborn, and RDKit libraries. We performed a one-way ANOVA followed by Tukey’s honestly significant difference (HSD) test to assess the statistical significance of differences in mean values.

### HTVS and enrichment factor calculation

All docking studies were performed using AutoDock Vina.^73^ Ligands were prepared with ‘scrub.py’ and ‘mk_prepare_ligand.py’, while proteins were prepared using ‘mk_prepare_protein.py’. HTVS was carried out with the default parameters of AutoDock Vina. For the Enamine HTS collection, about one-tenth of the molecules (∼178,000 compounds) were randomly chosen to balance chemical diversity with computational efficiency.

The enrichment factor^48^ is defined as follows:

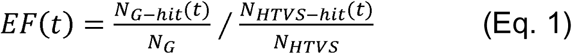

Where *N_G-hit_* (*t*) is the number of generated molecules with docking scores lower than defined docking score thresholds t. *N_G_* is the number of generated molecules. *N_HTVS-hit_* (*t*) is the number of HTVS molecules with docking scores lower than t, and *N_HTVS_* is the number of molecules in the HTVS library. If the EF value is greater than 1, it means the generative model has better effectiveness at finding molecules with higher affinity than the given docking score thresholds.

## Author Contributions

The manuscript was prepared with contributions from all authors who have approved the final version.

## Notes

The authors declare no competing financial interest.

## Acknowledgments

We sincerely thank Dr. Patrick Walters for his valuable suggestions and insights, which significantly improved this manuscript. We also extend our gratitude to Dr. Brian Shoichet (UC San Francisco), Dr. Jean-Louis Reymond (University of Bern), Dr. Conor Colley (Massachusetts Institute of Technology), Yu Chen (Freie Universität Berlin), and Ye Buehler (University of Bern) for their helpful discussions. This work was supported by NIH R01 awards (GM129431/GM143370) and the AnalytiXIN fellowship (to J.L.).

