## Supplemental information for "Structure-Based Generation of 3D Small-Molecule Drugs: Are We There Yet?"

**Table S1.** Summary of 3D small-molecule generative algorithms studied in this work. All the algorithms use Cartesian coordinates for 3D representations.

|  | **If one-shot generation** | **If include domain knowledge** | **GitHub Repo?** |
| --- | --- | --- | --- |
| 3D_generative_SBDD | $\boldsymbol{\times}$ | $\boldsymbol{\times}$ | https://github.com/luost26/3D-Generative-SBDD |
| pocket2mol | $\boldsymbol{\times}$ | $\boldsymbol{\times}$ | https://github.com/pengxingang/Pocket2Mol |
| Targetdiff | $\boldsymbol{\surd}$ | $\boldsymbol{\times}$ | https://github.com/guanjq/targetdiff |
| Decompdiff | $\boldsymbol{\surd}$ | $\boldsymbol{\surd}$ | https://github.com/bytedance/DecompDiff |
| Decompopt | $\boldsymbol{\surd}$ | $\boldsymbol{\surd}$ | https://github.com/bytedance/DecompOpt |
| Molsnapper | $\boldsymbol{\surd}$ | $\boldsymbol{\surd}$ | https://github.com/oxpig/MolSnapper |
| PMDM | $\boldsymbol{\surd}$ | $\boldsymbol{\times}$ | https://github.com/Layne-Huang/PMDM |

**Table S2.** Number of molecules generated by each model as well as three control groups. The number of valid molecules and the number of unique molecules based on the InChi key are also calculated. Some algorithms failed to generate 100 molecules for every receptor. We calculated the efficiency based on success cases.

|  | # of molecules | # of standardized molecules | # of unique molecules (InChi) | # of molecules have ring system | Efficiency (seconds per molecule) | GPU |
| --- | --- | --- | --- | --- | --- | --- |
| FDA approved small molecules | 2,778 | 2,777 | 2,590 | 2,281 (88%) | - | - |
| ChEMBL clinical small molecules | 8,236 | 8,233 | 7,488 | 7091 (95%) | - | - |
| CrossDocked2020 dataset | 11,759 | 11,596 | 11,314 | 10,638 (94%) | - | - |
| 3DSBDD | 10,499 | 9,955 | 9,803 | 8,952 (91%) | ~7.2 * | RTX 4070Ti Super |
| pocket2mol | 9,707 | 9,707 | 8,764 | 8,446 (96%) | ~16.2 | RTX 4070Ti Super |
| Targetdiff | 9,216 | 9,208 | 8,998 | 8,300 (92%) | ~15.5 | RTX 3080 |
| Decompdiff | 8,607 | 8,607 | 7,995 | 7,442 (93%) | ~21.0 | RTX 4070Ti Super |
| Decompopt | 8,913 | 8,913 | 8,280 | 7,604 (92%) | ~22.8 | RTX 3080 |
| Molsnapper | 5,606 | 5,606 | 5,589 | 5,302 (95%) | ~11.0 | RTX 4070Ti Super |
| PMDM | 9,757 | 9,612 | 9,263 | 9,099 (98%) | ~4.8 | RTX 3080 |

**Table S3.** Classification of targets from the Crossdock training and test set. The target classification follows BindingNetv2.^1^

| Protein Targets | Crossdock training set | Crossdock test set |
| --- | --- | --- |
| G-protein coupled receptor | 0 (0%) | 0 (0%) |
| Dehydratase | 22 (1.2%) | 0 (0%) |
| Esterase | 29 (1.6%) | 3 (3.2%) |
| Ion channel/transporter | 42 (2.3%) | 0 (0%) |
| Kinase | 168 (9.2%) | 13 (14.0%) |
| Nuclear receptor | 7 (0.4%) | 0 (0%) |
| Oxidoreductase | 245 (13.4%) | 19 (20.4%) |
| Phosphatase | 21 (1.8%) | 3 (3.2%) |
| Phosphodiesterase | 0 (0%) | 0 (0%) |
| Protease | 120 (6.6%) | 7 (7.5%) |
| Transferase | 347 (19.0%) | 18 (19.4%) |
| Transmembrane signal receptor | 0 (0%) | 0 (0%) |
| Others | 792 (43.4%) | 29 (31.2%) |
| **Total** | **1804** | **92** |

**Table S4**. Enrichment factors (EF) of generated molecules for each protein using various docking score cutoff values.

| Target | Redock score (kcal/mol) | Model | EF < redock | EF < -8 | EF < -9 | EF < -10 | EF < -11 | EF < -12 |
| --- | --- | --- | --- | --- | --- | --- | --- | --- |
| c-SRC kinase | -10.4 | 3DSBDD | 3.8 | 1.1 | 1.8 | 3.1 | 4.0 | 2.3 |
|  |  | pocket2mol | 10.7 | 1.4 | 2.5 | 7.0 | 19.2 | 39.3 |
|  |  | Molsnapper_np | 22.2 | 1.6 | 3.4 | 11.8 | 60.6 | 192.9 |
|  |  | Molsnapper_p | 13.0 | 1.5 | 2.7 | 7.7 | 29.9 | 77.2 |
| Smoothened | -11.3 | 3DSBDD | 6.1 | 0.7 | 0.8 | 1.5 | 4.5 | 10.9 |
|  |  | pocket2mol | 0 | 0.8 | 0.7 | 0.6 | 0.3 | 0 |
|  |  | Molsnapper_np | 17.8 | 1.3 | 2.0 | 5.4 | 18.1 | 10.7 |
|  |  | Molsnapper_p | 16.1 | 1.2 | 1.8 | 4.2 | 12.6 | 11.7 |
| Dopamine D1 receptor | -10.6 | 3DSBDD | 0 | 0.3 | 0.0 | 0.0 | 0.0 | 0.0 |
|  |  | pocket2mol | 12.9 | 1.2 | 2.0 | 4.9 | 22.5 | 1340.8 |
|  |  | Molsnapper_np | 18.0 | 1.1 | 2.5 | 11.2 | 10.9 | 366.3 |
|  |  | Molsnapper_p | 7.0 | 0.9 | 1.8 | 3.9 | 8.2 | 728.1 |

**Table S5.** PoseBuster^2^, PoseCheck^3^, and torsion angle analyses^4^ of 3D molecules generated by each algorithm on the CrossDock test set are summarized. Because several algorithms produced extreme outliers that led to unusually high average strain energies, the median strain energy was used as a more robust measure of central tendency. The best performance for each metric is shown in bold.

|  | Average passed PoseBuster tests per ligand (↑) | Average clashes per ligand (↓) | Median strain energy (kcal/mol) per ligand (↓) | Percentage strained torsion angle (↓) |
| --- | --- | --- | --- | --- |
| 3DSBDD | 20.1 / 21 | **5.7** | 334.7 | 60.9% |
| pocket2mol | 20.7 / 21 | 7.3 | 37.5 | **38.6%** |
| Targetdiff | 20.2 / 21 | 10.8 | 409.2 | 64.4% |
| Decompdiff | 20.5 / 21 | 10.5 | 167.7 | 68.8% |
| Decompopt | 20.5 / 21 | 10.1 | 177.2 | 70.4% |
| Molsnapper | 20.5 / 21 | 7.6 | 229.9 | 76.2% |
| PMDM | 19.3 / 21 | 23.4 | 91.1 | 41.8% |
| Crossdock test | **20.9 / 21** | 7.2 | **34.8** | 55.1% |

**Table S6.** The QED, SA score, and Retrosynthesis success rate (calculated using SynFormer)^5^ of molecules from three test sets and generated by different algorithms for the crossdock test set.

|  | QED (↑) | SA score (↓) | Retrosynthesis success rate (%)(↑) |
| --- | --- | --- | --- |
| FDA approved small molecules | 0.48 | 3.77 | 26.3% |
| ChEMBL clinical small molecules | 0.54 | 3.33 | 16.1% |
| CrossDocked2020 dataset | 0.52 | 3.34 | 15.7% |
| 3DSBDD | 0.50 | 4.46 | 6.8% |
| pocket2mol | 0.58 | 3.18 | 19.0% |
| Targetdiff | 0.49 | 4.61 | 5.7% |
| Decompdiff | 0.50 | 4.02 | 8.8% |
| Decompopt | 0.49 | 3.96 | 8.7% |
| Molsnapper | 0.39 | 4.11 | 4.2% |
| PMDM | 0.57 | 4.37 | 7.0% |

Besides the shared issues in our tested algorithms discussed in the main text, we analyzed each algorithm in more detail, focusing on specific aspects. For example,

- 3DSBDD and pocket2mol were designed to generate simple molecules, which could explain why pocket2mol performs better in the BM scaffolds metric we apply (Figures 4 and 5A), which could be a good thing if we could adopt the algorithm and use it for lead optimization or finding fragment hits.
- Molsnapper tends to generate more heteroatoms, averaging around 9.6 heteroatoms per molecule, which is significantly higher than those of any other method evaluated and control groups (Figure S5A). The structures shown in Figure S5B from Molsnapper raise concerns about chemical plausibility, synthetic accessibility, and metabolic liability due to the excess heteroatoms.
- PMDM prefers complex ring structures with fewer rotatable bonds (Figure S5C) with an average of fewer than 2 rotatable bonds per molecule, compared to approximately 5 in most other groups (excluding pocket2Mol and 3DSBDD). The scarce rotatable bonds from molecules generated by PMDM may correlate to its preference for bridgehead atoms (0.67 per molecule on average vs. less than 0.3 per molecule for control groups). The representative examples generated by PMDM (Figure S5D) further illustrate this tendency, showcasing structures that would be highly challenging to synthesize or unlikely to exist in practice.

For reference, we also compared the proposed metrics with QED, SA, and the retrosynthesis success rate predicted by a retrosynthesis model, as discussed here:

Quantitative Estimate of Drug-likeness (QED) and Synthetic Accessibility (SA) scores are two widely used metrics in the preclinical stage of drug discovery. QED is derived from the distribution of several properties across 771 approved orally available small molecules, producing a weighted sum score that estimates a compound’s potential to become an oral drug. In contrast, the SA score combines fragment frequency data from approximately one million representative compounds in PubChem with a structural complexity penalty to estimate the ease of synthesizing a molecule. However, SA does not penalize chemically implausible structures and is not designed to detect them. We calculated the average QED and SA scores for the three control groups as well as the generated molecules, and the results are summarized in Table S6. For QED, there is almost no distinction between the control sets and the generated molecules. For SA scores, although the control groups generally perform better than most generated molecules, the average scores of the generated molecules remain within an acceptable range (< 6). In addition, we recognize that current AI-based retrosynthesis models may help developers assess the synthetic accessibility of generated molecules by evaluating how many can be assigned a predicted synthetic route. We applied Synformer for this purpose, and the results are also shown in Table S6. The three control sets perform much better than the generated molecules. However, these retrosynthesis algorithms are themselves limited by generation efficiency (more than three days on one Nvidia 4070 Ti Super for ~10,000 molecules) and the coverage of their synthon libraries, and the proposed synthetic routes still require experimental validation.

In this study, we introduce alternative metrics that assess the occurrence of key ring structures or ring-plus-linker structures (i.e., BM scaffolds) in more drug-like datasets such as ChEMBL and subsets of ZINC. These metrics are intended to evaluate both chemical plausibility and drug-likeness of molecules generated by generative algorithms, with a stronger emphasis on identifying chemical plausible compounds (e.g., see the second structure in Figure 5B, SA score 4.1). Unlike QED and SA, these metrics serve a different purpose. We recommend applying these metrics to a sufficiently large set of generated molecules to gain an overall understanding of an algorithm’s performance in producing chemically plausible and drug-like compounds. They can also be used to filter and visualize specific molecules that may contain problematic ring structures or BM scaffolds. However, failing to meet these metrics does not necessarily imply that a specific molecule is chemically implausible or lacks drug-likeness.

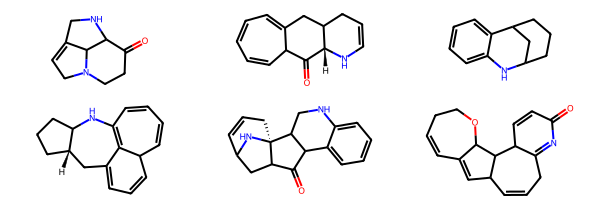

**Figure S1.** Examples of ring systems that appear zero time in ChEMBL database.

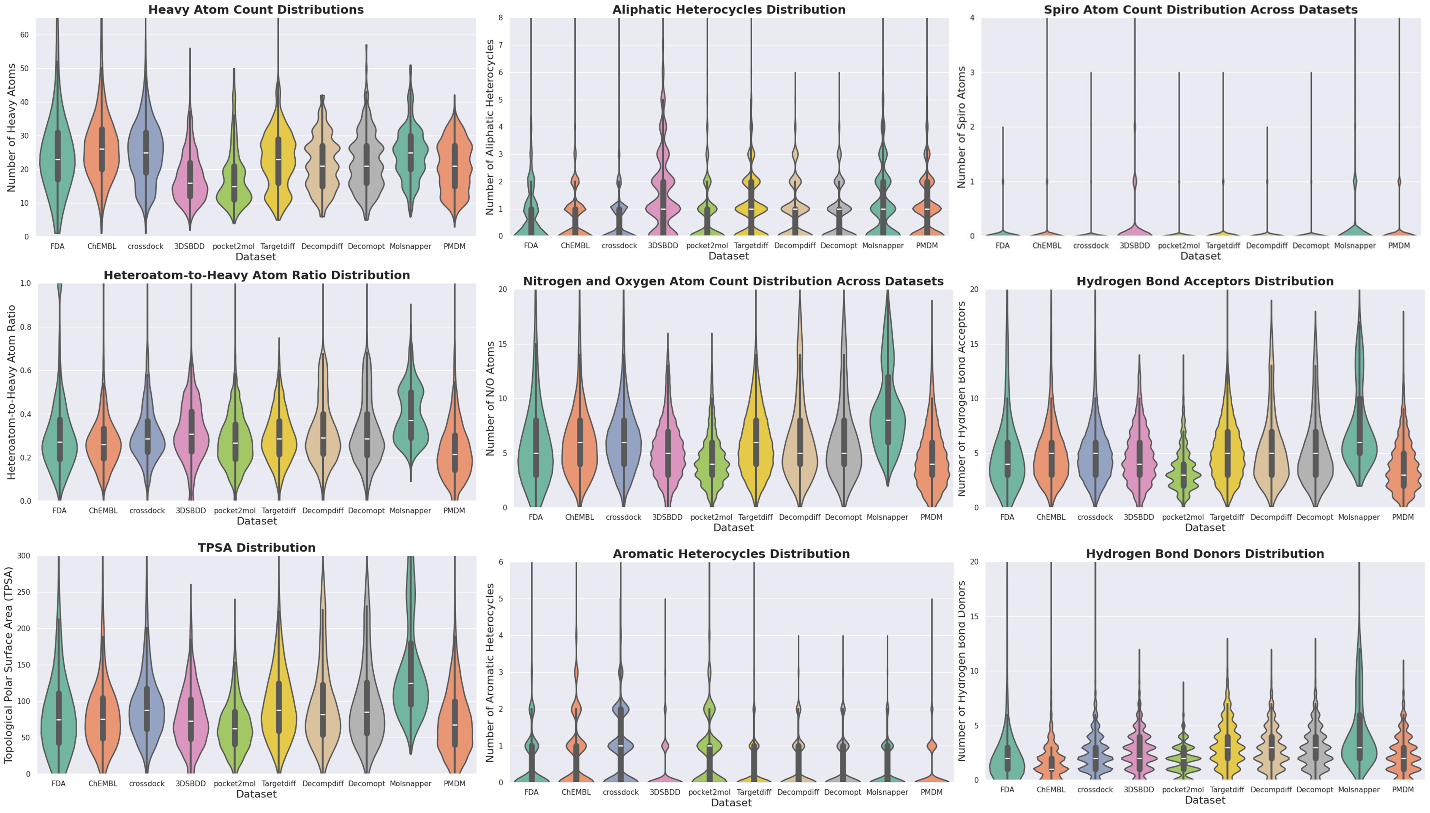

**Figure S2.** Other distributions calculated. The conclusion generally aligns with the main text, i.e., algorithms generate more aliphatic rings, fewer aromatic rings, more complex molecules, and favor polar functional groups. Overall, a detailed structural and property distribution analysis could provide more information about the potential weaknesses of algorithms and determine how to improve them, since not all chemically implausible molecules are flagged by the three metrics we used.

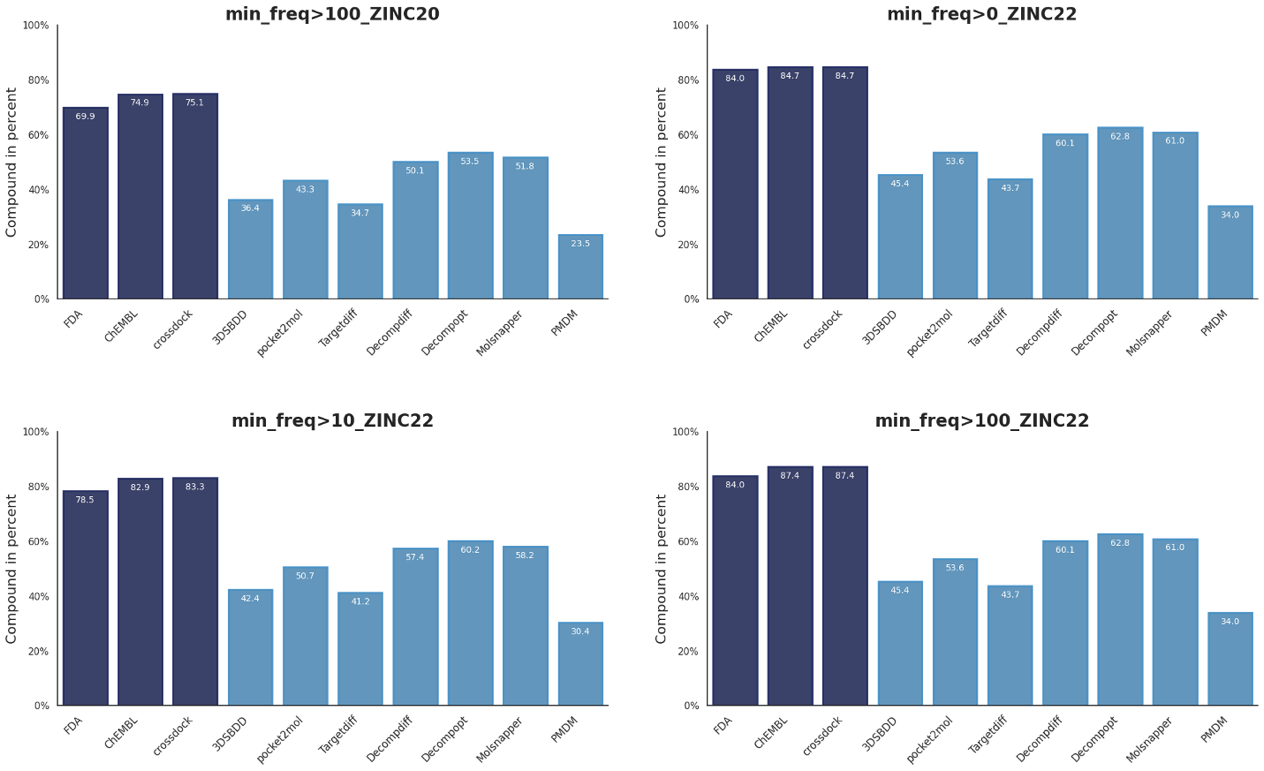

**Figure S3.** Benchmarking result by ring system metric using different thresholds and datasets. The conclusion remains consistent: the ring system metric could differentiate between control groups and molecules generated by algorithms.

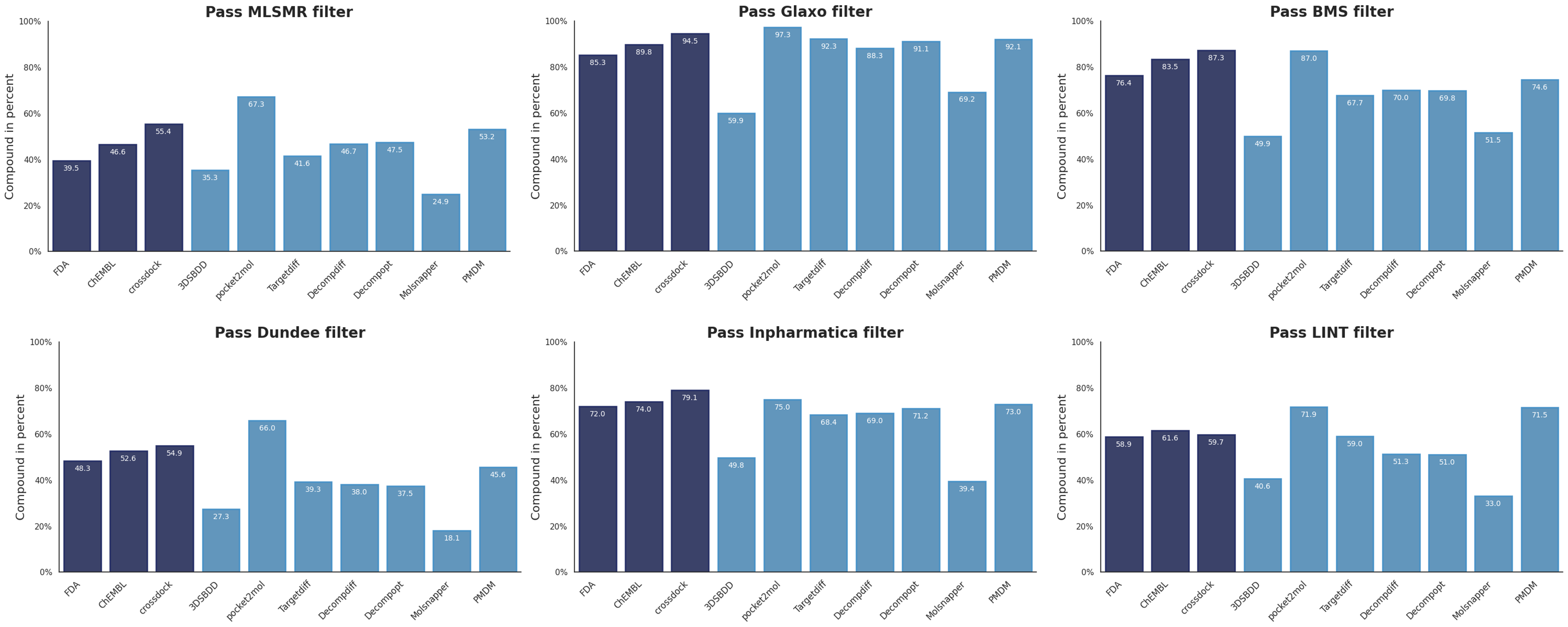

**Figure S4.** Commonly applied filters fail to differentiate between control groups and molecules generated by generative algorithms.

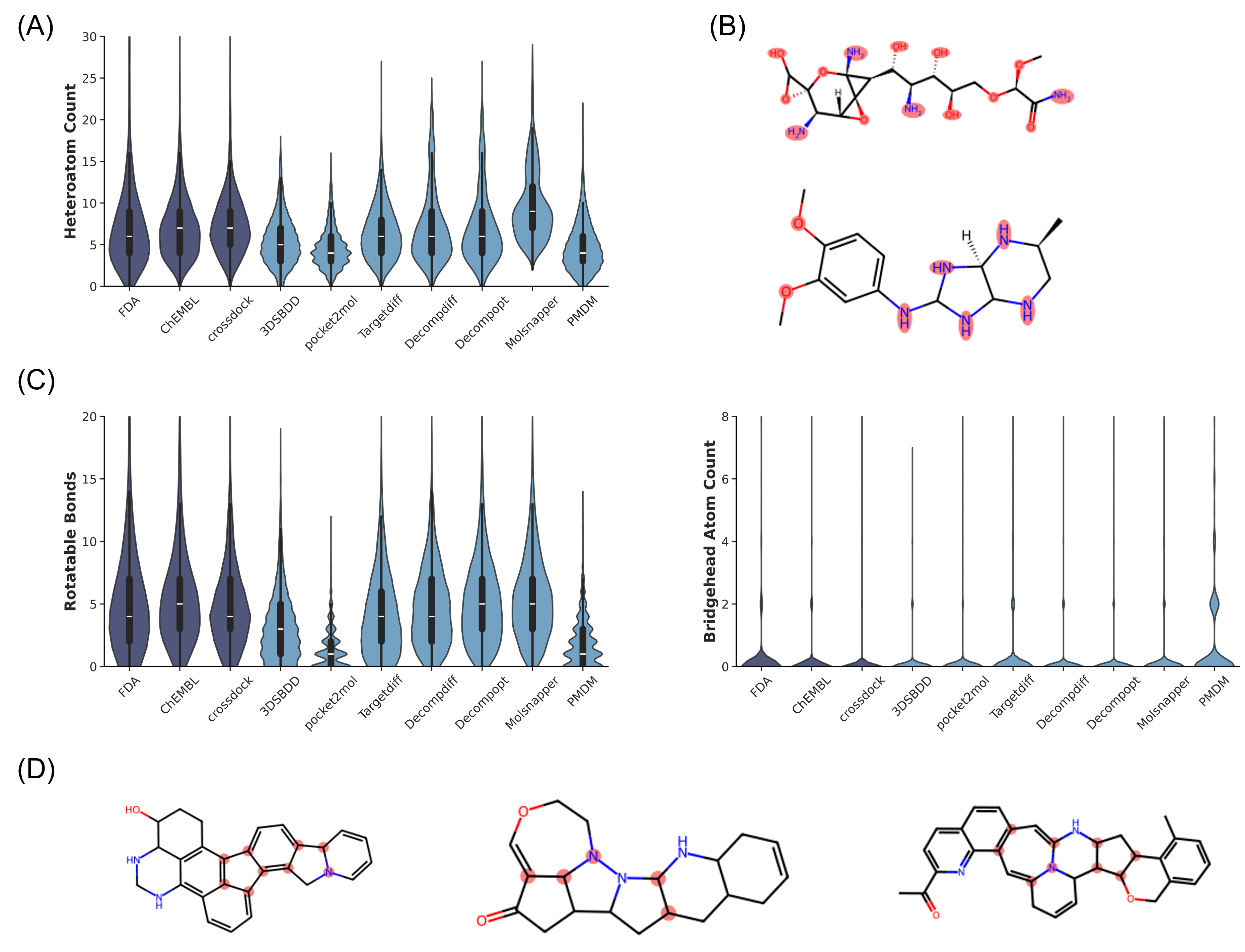

**Figure S5**. Distributions that provide specific information for single algorithm. Corresponding chemically implausible structures are selected from molecules that didn’t pass ChEMBL ring metrics and substructures associated with the distributions are highlighted in red. (A) Molsnapper tends to generate more heteroatoms than other algorithms and control datasets. (B) Example of molecules with multiple heteroatoms. All molecules are generated by the Molsnapper algorithm. (C) PMDM generates more bridged molecules with fewer rotatable bonds. (D) Example molecules with multiple bridgehead atoms. All molecules are generated by the PMDM algorithm.

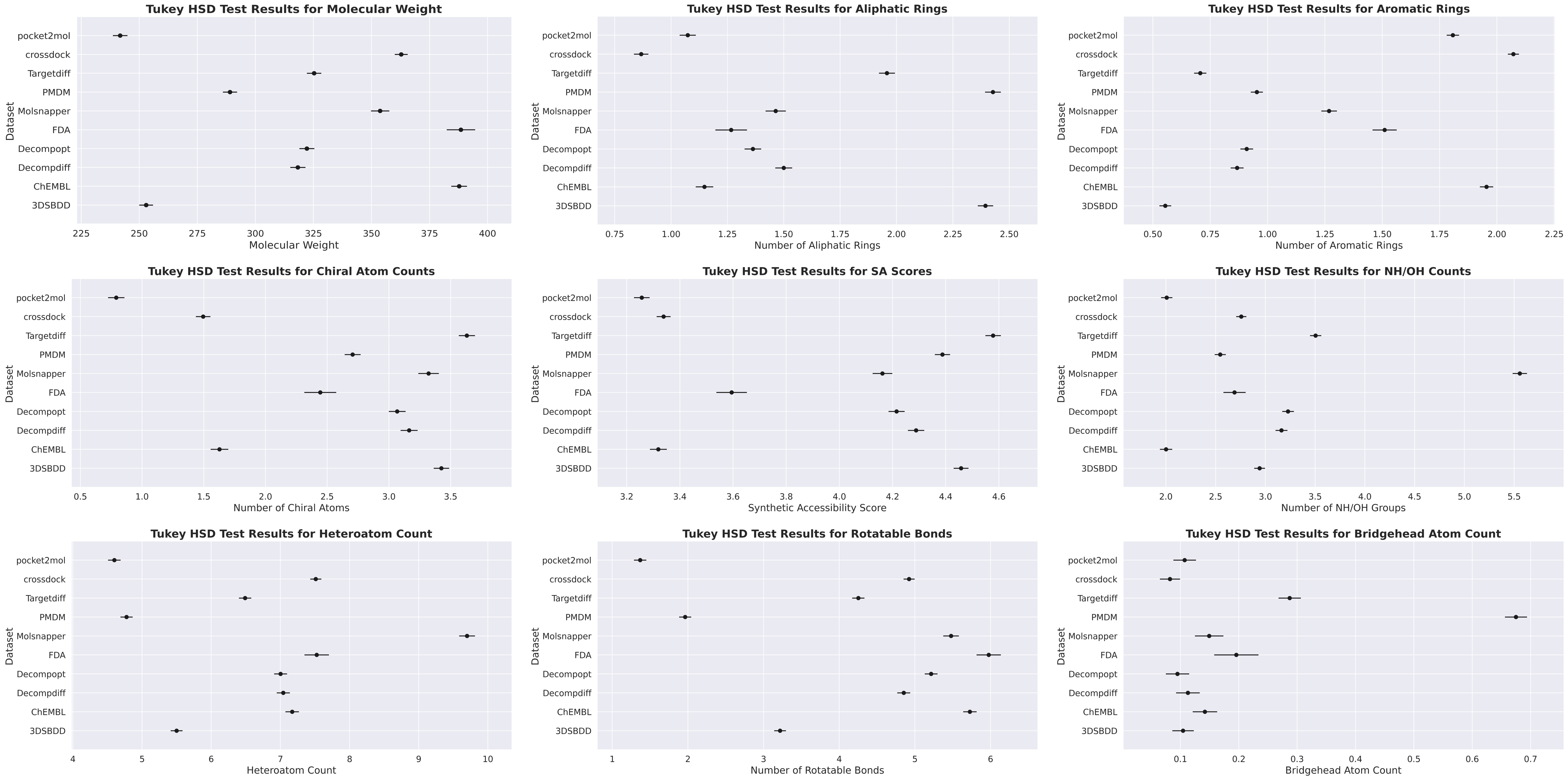

**Figure S6.** Results of Tukey’s honestly significant difference (HSD) test for each distribution plot, comparing the seven algorithms against the three control groups (no pairwise comparisons were listed here within the algorithm set or within the control set). Pairs listed below showed no significant difference (p > 0.05); all other algorithm–control comparisons were significantly different.: For aliphatic rings, Decompopt and FDA are not significantly different. For NH/OH count, ChEMBL and pocket2mol, FDA and PMDM are not significantly different. For heteroatom, ChEMBL and Decompdiff, ChEMBL and Decomopt are not significantly different. For rotatable bonds, CrossDock and Decompdiff are not significantly different. For bridgehead atom, 3DSBDD and ChEMBL, 3DSBDD and crossdock, ChEMBL and Decompdiff, ChEMBL and Molsnapper, ChEMBL and pocket2mol, Decompdiff and crossdock, Decompopt and crossdock, FDA and Molsnapper, crossdock and pocket2mol are not significantly different.

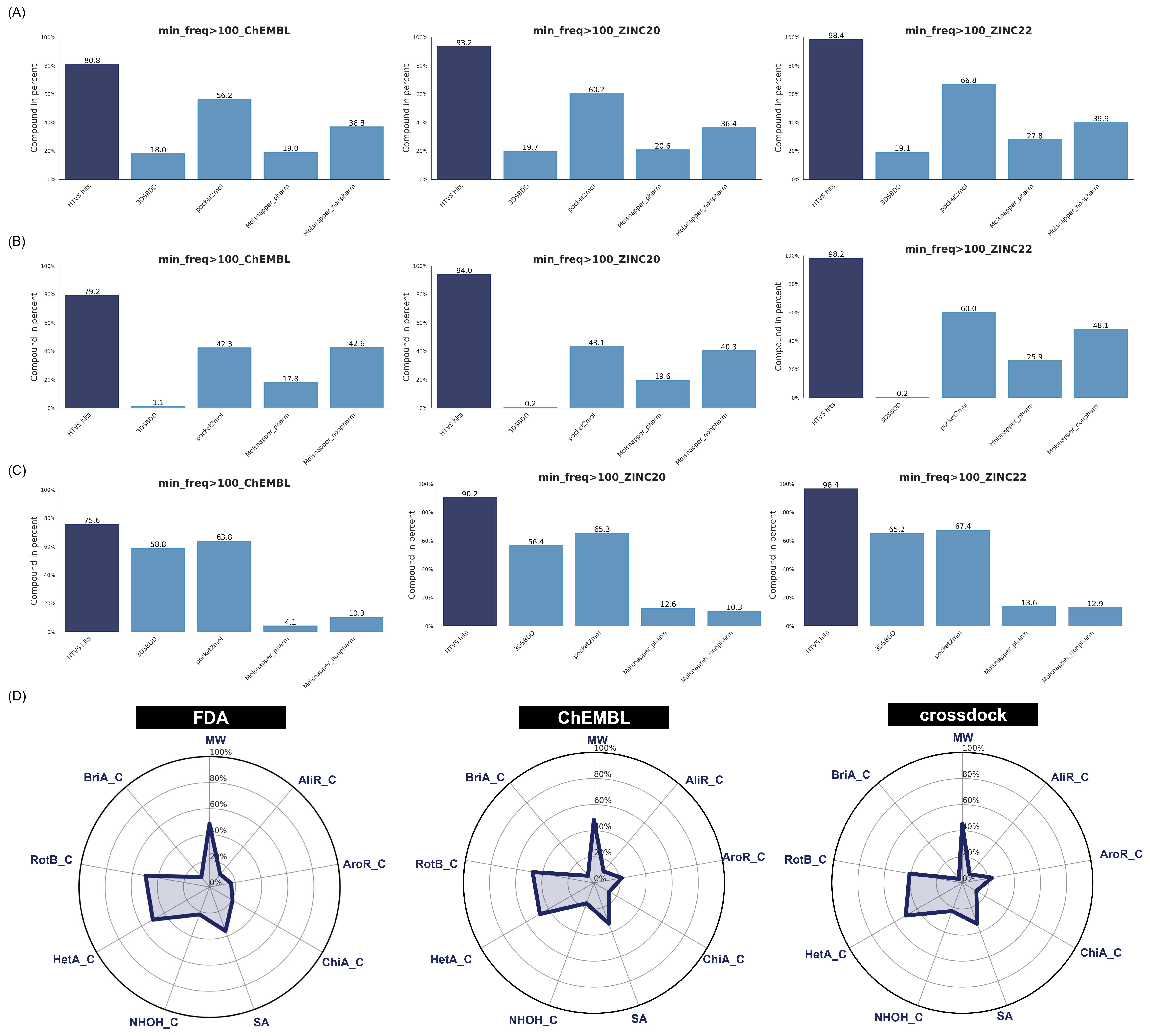

**Figure S7.** (A-C) Ring frequency metrics using ring systems extracted from different datasets. All metrics could distinguish between HTVS hits and generated molecules on three case study proteins. From top to bottom: c-Src kinase, Smoothened receptor and Dopamine D1 receptor. (D) Radar plot of the property distributions of three control groups. Percentages were calculated using the following values as denominators, with the average value serving as the numerator. The denominators used for normalization are same with Figure 8 for comparison. MW=800, AliR_C=10, AroR_C=9, ChiA_C=12, SA=10, NHOH_C=12, HetA_C=15, RotB_C=12, and BriA_C=2. The Enamine compounds may occupy relatively novel chemical space, making them difficult to capture with BM scaffold–based metrics but more readily characterized by ring frequency–based metrics.

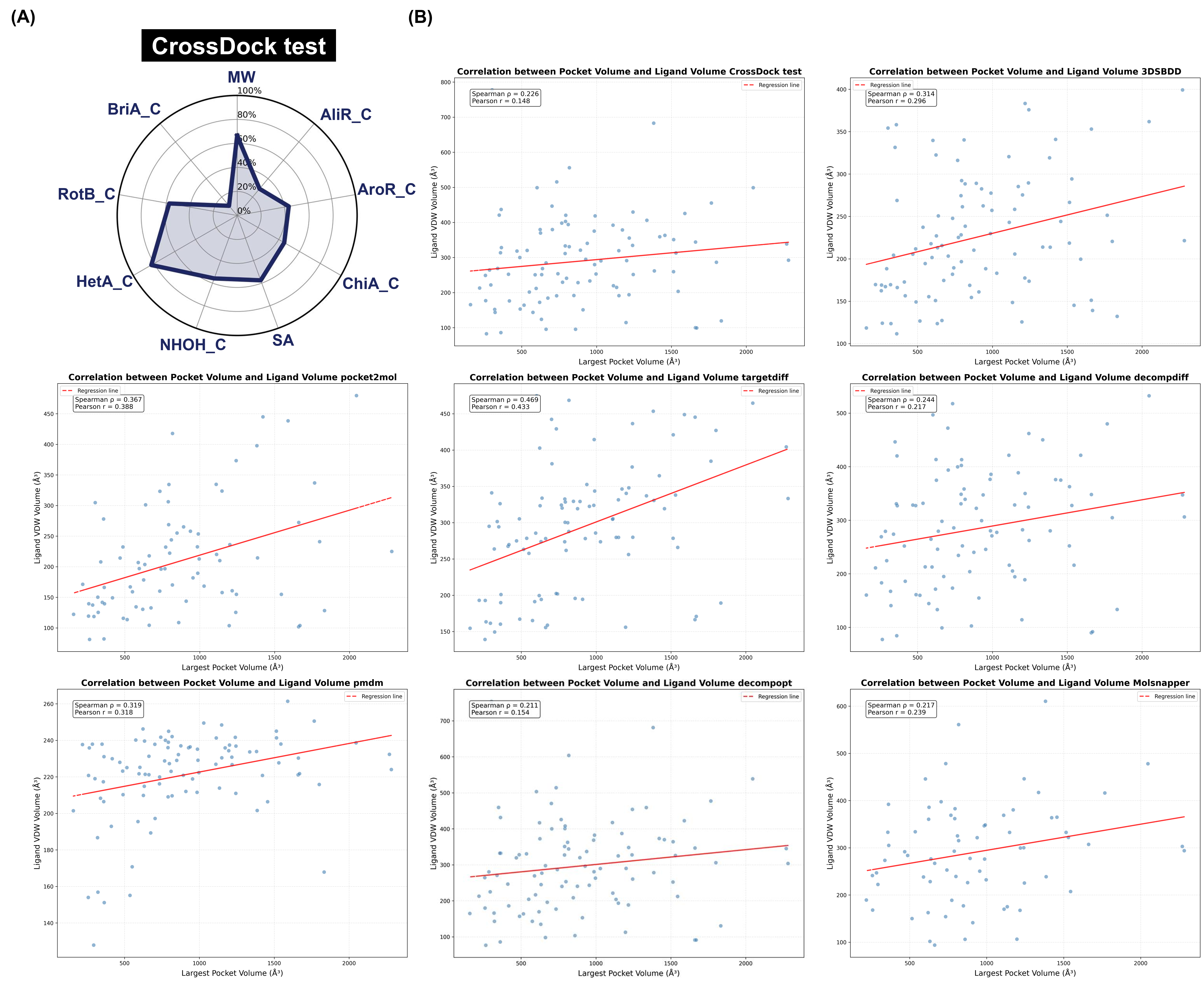

**Figure S8.** (A) Chemical property distribution radar plot of 100 ligands in crossdock test set. (B) The correlation between the volume of pocket from crossdock test set and the average van-der-Waals volume of generated ligands for this pocket. All algorithms and crossdock test sets show a weak positive correlation between pocket volume and ligand volume.
